# Performance Comparison of Computational Prediction Methods for the Function and Pathogenicity of Non-coding Variants

**DOI:** 10.1101/2021.10.05.463137

**Authors:** Zheng Wang, Guihu Zhao, Bin Li, Zhenghuan Fang, Qian Chen, Xiaomeng Wang, Tengfei Luo, Yijing Wang, Qiao Zhou, Kuokuo Li, Lu Xia, Yi Zhang, Xun Zhou, Hongxu Pan, Yuwen Zhao, Yige Wang, Lin Wang, Jifeng Guo, Beisha Tang, Kun Xia, Jinchen Li

## Abstract

Non-coding variants in the human genome greatly influence some traits and complex diseases by their own regulation and modification effects. Hence, an increasing number of computational methods are developed to predict the effects of variants in the human non-coding sequences. However, it is difficult for users with insufficient knowledge about the performances of computational methods to select appropriate computational methods from dozens of methods. In order to solve this problem, we assessed 12 performance measures of 24 methods on four independent non-coding variant benchmark datasets: (I) rare germline variant from ClinVar, (II) rare somatic variant from COSMIC, (III) common regulatory variant dataset, and (IV) disease associated common variant dataset. All 24 tested methods performed differently under various conditions, indicating that these methods have varying strengths and weaknesses under different scenarios. Importantly, the performance of existing methods was acceptable in the rare germline variant from ClinVar with area under curves (AUCs) of 0.4481 - 0.8033 and poor in the rare somatic variant from COSMIC (AUCs: 0.4984 - 0.7131), common regulatory variant dataset (AUCs: 0.4837 - 0.6472), and disease associated common variant dataset (AUCs: 0.4766 -0.5188). We also compared the prediction performance among 24 methods for non-coding *de novo* mutations in autism spectrum disorder and found that the CADD and CDTS methods showed better performance. Summarily, we assessed the performances of 24 computational methods under diverse scenarios, providing preliminary advice for proper tool selection and new method development in interpreting non-coding variants.

## Introduction

The most regions of human genome are non-coding sequences and some non-coding sequences harbours structural, regulatory, and transcribed information [1]. Some variants in the non-coding sequences play important roles in medical traits and complex diseases [2]. It is widely accepted that a large proportion of the non-coding sequences is functional and harbours genetic variants that contribute to disease aetiology [3] and that modified penetrance of pathogenic coding variants by non-coding regulatory variants can contribute to disease risk [4]. In addition, recent discoveries support that variants in non-coding sequences are important in cancer development [5, 6]. Furthermore, genome-wide association studies (GWASs) have identified numerous single-nucleotide variants (SNVs) associated with many medical traits and complex diseases, and most of these associations are thought to be mediated by non-coding regulatory variants [7–9].

In the last few years, many genomic features in the non-coding sequences of the genome have been identified across multiple human tissues and cell types through various large-scale projects, such as ENCODE [10], Roadmap Epigenomics [11] and FANTOM5 [12], enabling analysis and prediction of the functional effects of non-coding variants. Several computational methods [13–32] based on supervised, unsupervised, and semi-supervised models have been developed to prioritize non-coding variants by integrating various genomic features. For example, CADD used more than 60 various annotations from conservation, epigenetic modification, genetic context, and functional prediction [13]; PAFA was the first method to introduce the fixation index [33], a population level metric important for prioritizing population relevant functional non-coding variants [30]. Given that computational methods have varying advantages, disadvantages, and specific features [34], it is essential for different user requirements to choose appropriate methods. Three previous studies have evaluated the performance of several computational methods [35–37]. Nevertheless, limited benchmark datasets were used in the three studies, and they measured the area under the receiver operating characteristic (ROC) curves and area under the precision–recall (PR) curves; other critical performance measures, such as the accuracy at 95% sensitivity or specificity, were not used. Furthermore, several recently developed methods, such as ncER [28], DVAR [18], and PAFA [30], have not been evaluated in detail. Hence, it is particularly important to systematically and comprehensively evaluate these methods to help users to choose applicable computational methods.

Notably, in our previous research, we did not develop any computational method for non-coding variants. Therefore, we independently assessed 12 performance measures for 24 methods using four benchmark datasets. Our study compared computational methods under different conditions and showed that the performance of each method varied under different conditions. We also found some computational methods with an acceptable performance for rare pathogenic germline variants and noted that no methods had satisfactory prediction results for rare somatic, disease associated common, and common regulatory variants. Our results provide an opportunity for clinicians and researchers to select applicable evaluation methods to explore the functional effects of non-coding variants. Additional more accurate computational methods for various non-coding variants must be developed.

## Results

### Benchmarks showed poor concordance among the prediction scores of existing methods

In this study, a total of 24 computational methods was assessed (**Table 1**). Four independent benchmark datasets were built that represented various genetic aspects: the rare germline variant from ClinVar, which included rare non-coding germline variants of human medical traits and genetic diseases [38]; rare somatic variant from COSMIC for rare non-coding somatic variants of human cancers [39, 40]; common regulatory variant dataset for common non-coding variants of the human genome that explain variation in gene expression levels [41–43]and disease associated common variant dataset for common non-coding risk variants of human diseases that are recognized by GWASs [43, 44] (**Table 2**). Further, all 24 computational methods were published before 2020 and the training datasets used were published before 2019. To reduce overlap between our testing benchmark data and the training data used in the 24 computational methods, we selected variants published after 2019 and removed variants that existed in these publicly available training datasets before comparing the methods.

**Table 1.** Summary of 24 computational methods compared in this study.

| Method | Prediction model | Model type | Learning dataset | Version | Ref. |
| --- | --- | --- | --- | --- | --- |
| CADD | SVM and logistic regression model | Supervised | Simulated DNMs and variants arisen and fixed in human populations | v1.3 | [13] |
| CDTS | Difference between expected and observed score as context-dependent tolerance score | Unsupervised | 11,257 human whole-genome sequences | 2017 | [14] |
| CScape | Kernel-based models and leave-one-concentration-out cross validation | Supervised | Somatic point variants from the COSMIC and SNVs from the 1000 Genomes Project | 2017 | [15] |
| DANN | Deep neural network | Supervised | Simulated DNMs and variants arisen and fixed in human populations | 2015 | [16] |
| DIVAN_TSS | Ensemble learning framework | Supervised | Risk variants of 45 diseases/phenotypes (ARB) and benign variants are sampled from the 1000 Genomes Project with TSS-matched criterion | 2016 | [17] |
| DIVAN_REGION | Ensemble learning framework | Supervised | Risk variants of 45 diseases/phenotypes (ARB) and benign variants are sampled from the 1000 Genomes Project with region-matched criterion | 2016 | [17] |
| DVAR | Multivariate Dirichlet Process Mixtures | Unsupervised | 2 million variants randomly sampled from the 1000 Genomes Project | v1.0 | [18] |
| Eigen | Spectral meta-learner | Unsupervised | Variants in the 1000 Genomes Project without a match in dbNSFP and within 500 bp upstream of the TSS | v1.1 | [19] |
| Eigen_PC | Spectral meta-learner | Unsupervised | Variants in the 1000 Genomes Project without a match in dbNSFP and within 500 bp upstream of the TSS | v1.1 | [19] |
| FATHMM-MKL | Multiple kernel learning | Supervised | Germline variants in HGMD and control variants from the 1000 Genomes Project | 2017 | [20] |
| FATHMM-XF | Kernel-based models and platt scaling | Supervised | Positive variants from the HGMD and control variants from the 1000 Genomes Project | 2017 | [21] |
| FIRE | Random forest model | Supervised | cis-eQTL SNVs identified by the Geuvadis lymphoblastoid cell lines and sampled non-eQTL SNVs | 2017 | [22] |
| fitCons | Generative probabilistic model | Semi-supervised | Multiple species genomic DNA sequence | v1.01 | [23] |
| FitCons2 | Probabilistic evolutionary model | Semi-supervised | Multiple species genomic DNA sequence | 2017 | [24] |
| FunSeq2 | Weighted scoring scheme | Semi-supervised | Small-scale informative data context from the 1000 Genomes Project, ENCODE, COSMIC and CGC | v2.1.6 | [25] |
| GenoCanyon | Conditional joint density estimation | Unsupervised | Each location in the human genome | v1.0.3 | [26] |
| LINSIGHT | Combination of generalized linear model and probabilistic model | Semi-supervised | Multiple species genomic DNA sequence and 54 unrelated human genomes | 2017 | [27] |
| ncER | XGBoost model | Supervised | Positive examples from HGMD (2016_R1) and ClinVar (July 2016) and negative examples from gnomAD | v1.0 | [28] |
| Orion | Difference between the observed and expected site-frequency spectrums | Unsupervised | 1662 WGS samples | 2017 | [29] |
| PAFA | Logistic regression with L1 regularization | Supervised | Variants labeled “pathogenic” in ClinVar and significant SNPs associated with complex traits or diseases and variants labeled “benign” in ClinVar and variants in the 1000 Genomes Project | 2018 | [30] |
| regBase_REG | XGBoost model | Supervised | Functional regulatory variants dataset and non-coding variants from the 1000 Genomes Project | v1.0 | [31] |
| regBase_PAT | XGBoost model | Supervised | Pathogenic regulatory variants dataset and non-coding benign variants labeled “benign” in ClinVar | v1.0 | [31] |
| regBase_CAN | XGBoost model | Supervised | Cancer recurrent regulatory somatic mutations dataset and non-recurrent somatic mutations | v1.0 | [31] |
| ReMM | Random forest model | Supervised | Hand-curated set of regulatory Mendelian mutations and derived alleles of human evolution | v0.3.1 | [32] |

*Note:* The difference between DIVAN\_TSS and DIVAN\_REGION was criteria to choose benign variants in the training set. Eigen\_PC had the same prediction model and learning dataset as Eigen but they had different weights for some genomic features. regBase trained three composite models based on different training datasets to score functional, pathogenic and cancer driver non-coding regulatory variants respectively. *de novo* mutations, DNMs; COSMIC, the catalogue of somatic mutations in cancer; SNVs, single nucleotide variants; ARB, association results browser; TSS, transcription start site; HGMD, human gene mutation database; eQTL, expression quantitative trait loci; ENCODE, Encyclopedia of DNA Elements; CGC, cancer gene census; WGS, Whole Genome Sequencing.

**Table 2.** Summary of four independent benchmark datasets used in this study.

| Benchmark dataset | Positive set | Negative set | Number of positive variants | Number of negative variants | Ref. |
| --- | --- | --- | --- | --- | --- |
| Rare germline variant from ClinVar | Non-coding ‘pathogenic’ and ‘likely pathogenic’ germline variants from ClinVar (20190102 -20201128) | Non-coding ‘benign’ germline variants from ClinVar (20190102 -20201128) | 515 | 1850 | [38] |
| Rare somatic variant from COSMIC | Non-coding somatic variants from COSMIC (v88 - v92) with recurrence $\geq 2$ and located on risk genes collected by CNCD | Non-coding somatic variants from COSMIC (v88 -v92) with recurrence = 1 and located genes except for risk genes collected by CNCD | 1966 | 597,221 | [39, 40] |
| Common regulatory variant dataset | eQTL SNPs from the GTEx Portal database and Brown's study | Randomly selecting variants with matched properties<sup>a</sup> from the 1000 Genomes Project by vSampler | 13,274 | 13,274 | [41-43] |
| Disease associated common variant dataset | Non-coding SNVs in the intersection set of credible sets defined by three tools<sup>b</sup> from CAUSALdb database with MAFs > 5% | Non-coding SNVs from the 1000 Genomes Project with MAFs > 5% in the same LD blocks as corresponding positive variants with $r^2$ thresholds > 0.2 | 73,693 | 76,214 | [43, 44] |

*Note:* <sup>a</sup> Matched properties including minor allele frequency, distance to closet transcription start site, gene density and number of variants in linkage disequilibrium.
<sup>b</sup> Three tools including PAINTOR [62], CAVIARBF [63], and FINEMAP [64]. COSMIC, the catalogue of somatic mutations in cancer; CNCD, Cornell Non-coding cancer driver database; eQTL, expression quantitative trait loci; GTEx, genotype-tissue expression; MAFs, major allele frequencies; LD, linkage disequilibrium; SNPs, single nucleotide polymorphisms; SNVs, single nucleotide variants.

Spearman rank correlation coefficients were calculated between any two computational methods based on the PHRED-scaled scores of four benchmark datasets to evaluate the predictive concordances among the 24 computational methods (**Figure S1**). The overall pairwise correlation for the rare somatic variant from COSMIC was generally higher than that for the other three datasets, suggesting that current methods show better concordance in somatic variant prediction. Moreover, we calculated the Spearman rank correlation coefficient based on the positive variant dataset and negative variant dataset for each benchmark dataset and found that the overall pairwise correlation for the negative rare somatic variant from COSMIC was higher than that for the positive rare somatic variant from COSMIC. The weak pairwise correlations (R < 0.4) among all 24 computational methods were common in the four benchmark datasets, except that a few computational methods were highly correlated with each other (R > 0.8) in the positive rare germline variant from ClinVar, such as CADD and DANN, possibly because of the selection of similar training data and learning features. In summary, our results indicate that existing computational methods have poor predictive concordance for the same benchmark dataset, suggesting the necessity and importance of assessing different computational methods under various conditions.

### Existing methods showed different performance for rare germline and rare somatic variants

It is widely accepted that pathogenic variants are often rare variants. To determine the performance of all 24 methods for rare variants, we constructed two datasets including rare germline variant from ClinVar and rare somatic variant from COSMIC. (□) The rare germline variant from ClinVar included 515 positive and 1850 negative variants (Table 2 and Table S1), which were downloaded from ‘pathogenic’, ‘likely pathogenic’, and ‘benign’ non-coding germline variants in the ClinVar database [38] with allele frequencies (AFs) < 0.1% in the GnomAD database [45]. (□) The rare somatic variant from COSMIC included 1966 positive and 597,221 negative variants (Table 2 and Table S1), and all of these variants were downloaded from the COSMIC database [39] with AFs < 0.1% in the GnomAD database. In addition, we selected the AUC as our major performance measure because its value is not affected by different cut-off values compared to other measures.

Assessments of 12 performance measures for all 24 computational methods based on the PHRED-scaled scores of the rare germline variant from ClinVar are summarised in **Table 3**. We found that the AUCs of the 24 methods ranged from 0.4481 to 0.8033 (median, 0.6988), and that FATHMM-XF (AUC = 0.8033) exhibited the best performance, followed closely by FATHMM-MKL (AUC = 0.7954) and ReMM (AUC = 0.7848). Because clinicians and researchers sometimes require computational methods with high sensitivity or specificity (typically > 95%) and they may choose computational methods to evaluate the pathogenicity of non-coding variants in genetic counselling for known pathogenic genes, which would be expected to identify disease-causing variants with a high sensitivity (true-positive rate > 95%), we further assessed the high-specificity regional (hspr)-AUC and high-sensitivity regional (hser)-AUC values. We found that FATHMM-XF (hspr-AUC = 0.7067) exhibited the best performance with hspr-AUC values > 0.70; regBase_PAT (hser-AUC = 0.5517) exhibited the best performance with hser-AUC values > 0.55 (Table 3). The accuracy and Mathews correlation coefficient (MCC) were also used to assess the performance of computational methods, with FATHMM-XF showing the highest accuracy and MCC scores among the 24 methods. Notably, methods based on supervised models (median of AUCs = 0.7161) showed better performance than those based on semi-supervised models (0.6832) and methods based on unsupervised models (0.5961). Moreover, we assessed the performance of the 24 computational methods based on the rare germline variant from ClinVar after removing the ‘likely pathogenic’ non-coding germline variants, resulting in 343 positive variants and 1850 negative variants. The results of assessment of 12 performance measures for all 24 computational methods are summarised in Table S2. Performance measures such as the AUCs of the computational method were generally concordant, regardless of whether the variants were likely pathogenic (Figure S2).

**Table 3.** Performance evaluation based on the rare germline variant from ClinVar.

| Method | Missing (%) | Best-threshold | PPV (%) | NPV (%) | FNR (%) | Sensitivity (%) | FPR (%) | Specificity (%) | Accuracy (%) | MCC | AUC | hspr-AUC | hser-AUC | Prediction model |
| --- | --- | --- | --- | --- | --- | --- | --- | --- | --- | --- | --- | --- | --- | --- |
| CADD | 0.00 | 11.1395 | 44.40 | 87.83 | 39.22 | 60.78 | 21.19 | 78.81 | 74.88 | 0.3572 | 0.7509 | 0.5587 | 0.5277 | SM |
| CScape | 9.77 | 30.6855 | 37.46 | 85.79 | 48.43 | 51.57 | 22.75 | 77.25 | 71.88 | 0.2589 | 0.6655 | 0.5344 | 0.5217 | SM |
| DANN | 0.00 | 9.5376 | 45.95 | 86.78 | 44.85 | 55.15 | 18.05 | 81.95 | 76.11 | 0.3484 | 0.7341 | 0.5956 | 0.5244 | SM |
| DIVAN_REGION | 0.00 | 2.6953 | 23.89 | 82.34 | 27.57 | 72.43 | 64.22 | 35.78 | 43.76 | 0.0715 | 0.5357 | 0.5153 | 0.5064 | SM |
| DIVAN_TSS | 0.00 | 3.8817 | 23.16 | 80.52 | 33.59 | 66.41 | 61.35 | 38.65 | 44.69 | 0.0431 | 0.5047 | 0.5040 | 0.5028 | SM |
| FATHMM-MKL | 0.00 | 12.1444 | 49.10 | 90.16 | 31.46 | 68.54 | 19.78 | 80.22 | 77.67 | **0.4375** | **0.7954** | **0.6359** | 0.5344 | SM |
| FATHMM-XF | 9.77 | 26.0395 | **60.32** | **90.98** | 33.18 | 66.82 | **11.61** | **88.39** | **83.88** | **0.5322** | **0.8033** | **0.7067** | 0.5074 | SM |
| FIRE | 0.00 | 9.7388 | 26.09 | 80.95 | 53.59 | 46.41 | 36.59 | 63.41 | 59.70 | 0.0831 | 0.5256 | NA | 0.5034 | SM |
| ncER | 0.25 | 13.9272 | 39.92 | 87.50 | 37.94 | 62.06 | 26.02 | 73.98 | 71.39 | 0.3144 | 0.7067 | 0.5249 | 0.5161 | SM |
| PAFA | 8.03 | 1.1395 | 36.40 | 90.42 | 26.32 | 73.68 | 34.15 | 65.85 | 67.49 | 0.3256 | 0.7239 | 0.5208 | NA | SM |
| regBase_CAN | 0.00 | 10.2935 | 39.50 | 89.49 | 29.51 | 70.49 | 30.05 | 69.95 | 70.06 | 0.3423 | 0.7083 | 0.5176 | 0.5018 | SM |
| regBase_PAT | 0.00 | 7.3824 | 35.23 | 89.40 | 26.60 | 73.40 | 37.57 | 62.43 | 64.82 | 0.2970 | 0.7375 | 0.5721 | **0.5517** | SM |
| regBase_REG | 0.00 | 20.6843 | 27.53 | 80.37 | 65.63 | 34.37 | 25.19 | 74.81 | 66.00 | 0.0852 | 0.5491 | NA | NA | SM |
| ReMM | 0.00 | 13.8161 | 47.68 | 89.36 | 34.17 | 65.83 | 20.11 | 79.89 | 76.83 | 0.4115 | **0.7848** | 0.5969 | **0.5448** | SM |
| CDTS | 8.25 | 10.9826 | 25.36 | 79.56 | 66.88 | 33.12 | 27.24 | 72.76 | 64.10 | 0.0538 | 0.4910 | NA | NA | UM |
| DVAR | 0.00 | 15.0531 | **51.63** | 88.69 | 38.45 | 61.55 | **16.05** | **83.95** | **79.07** | 0.4283 | 0.7420 | 0.5371 | 0.5159 | UM |
| Eigen | 10.40 | 15.3901 | 43.68 | 89.58 | 35.48 | 64.52 | 21.42 | 78.58 | 75.70 | 0.3786 | 0.7656 | 0.5379 | **0.5425** | UM |
| Eigen_PC | 10.40 | 8.8690 | 27.96 | 88.79 | **24.42** | **75.58** | 50.15 | 49.85 | 55.12 | 0.2064 | 0.6032 | NA | 0.5366 | UM |
| GenoCanyon | 0.00 | 12.2531 | 33.39 | 82.57 | 58.25 | 41.75 | 23.19 | 76.81 | 69.18 | 0.1721 | 0.5890 | 0.5418 | 0.5012 | UM |
| Orion | 14.80 | 11.1831 | 23.61 | 80.75 | 67.07 | 32.93 | 27.47 | 72.53 | 64.42 | 0.0488 | 0.5124 | 0.5074 | NA | UM |
| fitCons | 8.16 | 0.2856 | 21.12 | **95.45** | **0.22** | **99.78** | 98.78 | 1.22 | 21.87 | 0.0408 | 0.4481 | NA | 0.5034 | SSM |
| FitCons2 | 8.03 | 17.3066 | 41.81 | 86.30 | 48.46 | 51.54 | 19.02 | 80.98 | 74.80 | 0.3023 | 0.6909 | 0.6052 | 0.5029 | SSM |
| FunSeq2 | 1.61 | 7.6079 | 29.59 | **91.72** | **14.03** | **85.97** | 56.84 | 43.16 | 52.47 | 0.2491 | 0.6756 | 0.5330 | 0.5299 | SSM |
| LINSIGHT | 1.82 | 17.6486 | **64.97** | 88.15 | 44.44 | 55.56 | **8.31** | **91.69** | **83.85** | **0.5010** | 0.7743 | **0.6307** | 0.5005 | SSM |

*Note:* Best-threshold, the threshold corresponding to the best sum of sensitivity and specificity; PPV, positive predictive value; NPV, negative predictive value; FPR, false positive rate; FNR, false negative rate; MCC, mathew correlation coefficient; AUC, area under the curve; hspr-threshold, high-specificity regional threshold; hspr-AUC, high-specificity regional area under the curve; hser-threshold, high-sensitivity regional threshold; hser-AUC, high-sensitivity regional area under the curve; NA, not available; SM, supervised model; UM, unsupervised model; SSM, semi-supervised model. Top three methods of every measure are represented by bold text.

In addition, we assessed the performance of 24 methods for somatic variants and assessments of 12 performance measures based on PHRED-scaled scores, as summarised in Table S3. The AUCs of the 24 computational methods ranged from 0.4984 to 0.7131 (median, 0.6295) in the rare somatic variant from COSMIC, with FunSeq2 (AUC = 0.7131) exhibiting the best overall performance, followed closely by FitCons2 (0.7069). This result suggests that existing methods perform poorly for non-coding somatic variants. Furthermore, methods based on semi-supervised models (median of AUCs = 0.6988) performed better than methods based on unsupervised (0.6551) and supervised (0.6063) models.

### Predictive ability of existing methods for common variants should be improved

It is now accepted that some common variants are regulatory or risk variants; hence, we also constructed the common regulatory variant dataset and disease associated common variant datasets (see the MATERIALS AND METHODS section) to evaluate the performance of 24 methods for variants in the 1000 Genomes Project [43] with AFs > 5% (Table 2 and Table S1). The number of positive and negative variants, respectively, were recorded in the common regulatory variant dataset (13,274 and 13,274) and disease associated common variant dataset (73,693 and 76,214). We found that the AUCs of the 24 computational methods ranged from 0.4837 to 0.6472 (median, 0.5619) in the common regulatory variant dataset and from 0.4766 to 0.5188 (median, 0.5041) in the disease associated common variant dataset (Tables S4 and S5) and that the distributions of PHRED-scaled scores for positive and negative variants were similar whether they were in the common regulatory variant dataset or disease associated common variant dataset (Figures S3 and S4). This indicates that existing methods are not suitable for common variants, particularly for common variants in the same linkage disequilibrium (LD) block. Furthermore, we classified the disease associated common variant dataset into four subgroups (0.2 – 0.4, 0.4 – 0.6, 0.6 – 0.8 and 0.8 – 1.0) according to r^2^ thresholds of LD to observe the performance of all methods and found that all methods showed poor performance for four subgroups (Figure S5).

### CADD and CDTS showed better performance for non-coding *de novo* mutations (DNMs) in autism spectrum disorder (ASD)

Non-coding DNMs play important roles in neurodevelopmental disorders [46] such as DNMs in the promoter and regulatory regions in ASD [47, 48]. We then downloaded 115,569 and 113,530 non-coding DNMs from 1902 patients with ASD and 1902 unaffected siblings from the Gene4Denovo database [49] and evaluated the performance of the methods based on their PHRED-scaled scores (**Figure 1**, Table S6). Given that the pathogenicity of most non-coding DNMs is not clear, we selected odds ratios (ORs) to assess the performance of the computational methods; better methods were expected to show higher ORs under the same conditions. We adopted two strategies to calculate the ORs and two-sided *P* values between patients with ASD and their unaffected sibling.

**Figure 1.**
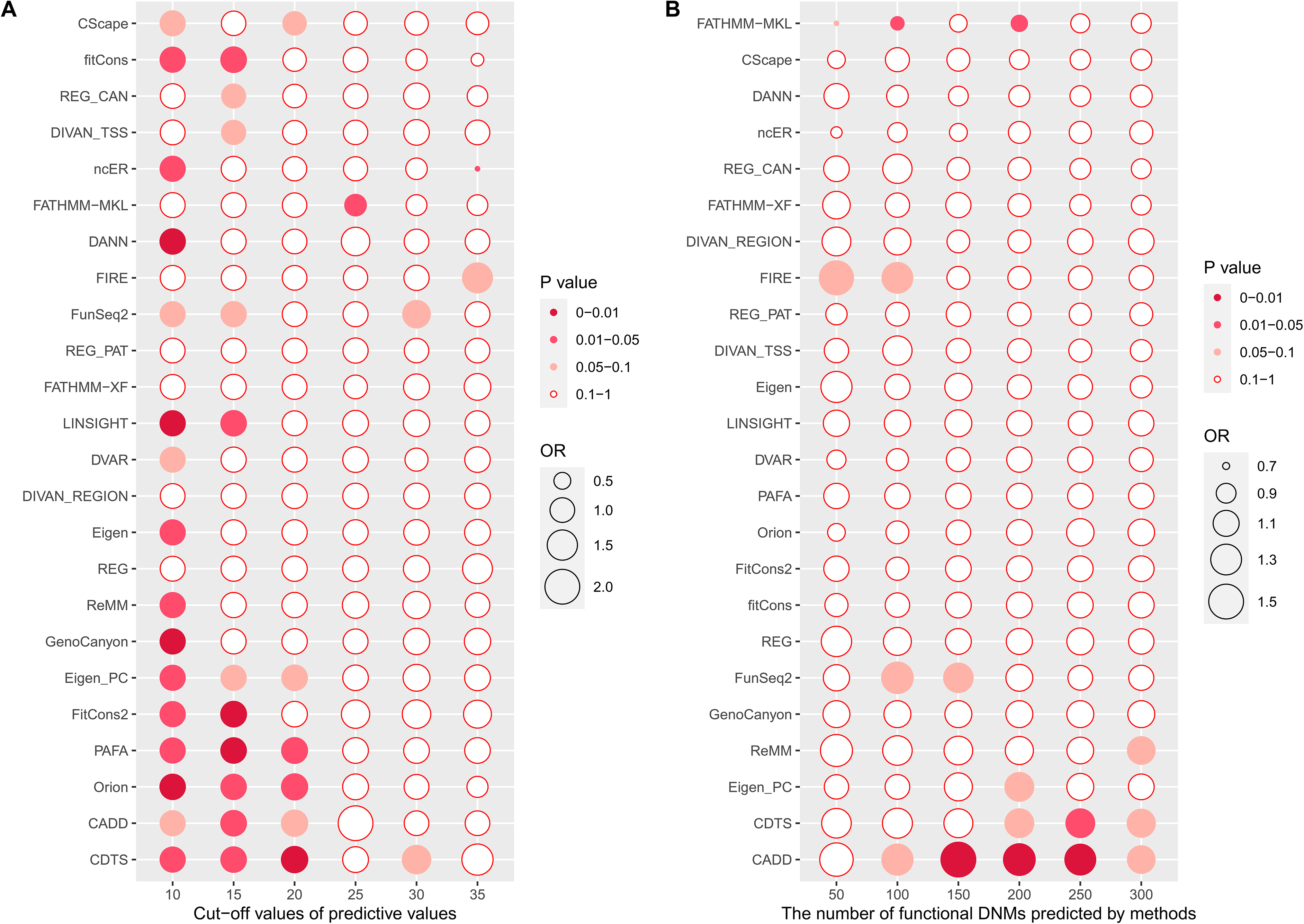
Performance evaluation based on *de novo* mutations (DNMs) **A**. Performance evaluation under different cut-off values. The x-axis represents different cut-off values of computational methods and y-axis represents different computational methods. The order of the computational methods is based on the size of their odds ratios (ORs) (cut-off value = 20). **B**. Performance evaluation under a different number of DNMs that are most likely to be functional in autism spectrum disorder (ASD). The x-axis represents the number of DNMs that are potentially functional variants in ASD as predicted by computational methods and y-axis represents different computational methods. The order of computational methods is based on the size of ORs (top 200). The OR and *P* values were calculated by two-sided Poisson’s ratio test. The area of each ball is proportional to the ORs. Different coloured balls represent different *P* value ranges.

In the first strategy, we counted the number of positive non-coding DNMs in the ASD and sibling groups under different cut-off values of PHRED-scaled scores (i.e., 10, 15, 20, 25, 30, and 35) for the 24 computational methods. The number of positive DNMs predicted by most methods between the ASD and sibling groups showed significant differences (*P* < 0.05) under the most relaxed condition (threshold = 10) but had low ORs (OR < 1.05). Under increasingly rigorous thresholds, many methods showed higher ORs but with *P* values > 0.05; the CDTS method achieved the best performance at a cut-off value of 20 (OR = 1.13, *P* = 0.006).

In the second strategy, we selected the top 50, 100, 150, 200, 250, and 300 DNMs that were most likely to be functional in patients with ASD based on PHRED-scaled scores and obtained corresponding thresholds to make predictions in unaffected siblings. We found that many methods yielded *P* values > 0.05 and ORs > 1.05 under the most relaxed condition (top 300). Under a more rigorous condition, some methods exhibited higher ORs and lower *P* values; CADD achieved the highest OR and lowest *P* value (OR = 1.5, P = 0.002, threshold = 21.6241), followed by CDTS (OR = 1.21, *P* = 0.0493, threshold = 26.8855). In summary, these results suggest that CADD and CDTS have better prediction performance for functional DNMs.

### Different methods showed different resolutions

Theoretically, a perfect computational method should assign different prediction scores to different variants at the same position. Here, we calculated the rates of discriminable prediction scores among 24 computational methods for the same position based on the whole genome and noted that only nine methods, including regBase_REG, regBase_CAN, and regBase_PAT, showed discriminability at base-wise resolution for most sites in the whole genome (Figure S6). Additionally, for computational methods without discriminability at base-wise resolution, we calculated the physical distances of surrounding DNA sites that showed the same prediction scores. We also determined the cumulative sum of proportions of different physical distances from 1 to the largest value until it was not smaller than 0.9, and then selected the last physical distances as the resolution. We found that most prediction scores of DNA sites differed with 1 bp site around them (Figure S7).

## Discussion

In recent years, it has been widely accepted that non-coding variants play important roles in human diseases [2–9]. Many computational methods for evaluating the function and pathogenicity of non-coding variants have been developed for clinicians and geneticists to help them identify functional or pathogenic non-coding variants. Given that computational methods for non-coding variants have adopted various algorithms and training data based on different evolutionary constraints, epigenomics, and sequence features, their performance differs under differing conditions. However, it is difficult to choose an optimal method because of the lack of knowledge about the performance of the methods under different conditions. Selecting an optimal method can effectively aid in the prioritization of functional variants and candidate genes, thus, increasing the demand for assessment of different computational methods under various conditions. In this work, we assessed 12 performance measures of 24 computational methods based on four non-coding independent benchmark datasets.

Although multiple studies [35–37] have compared computational prediction methods for non-coding variants, our study differs from these studies for the following reasons. (□) Our benchmark data are more comprehensive and strictness. We constructed four benchmark datasets representing different genomic contexts and simulated realistic situations, such as positive and negative variants from the common regulatory variant dataset with matched genomic features. (□) Our evaluation measures are more comprehensive. We not only selected some classic measures, but also adopted high-sensitivity regional AUC and high-specificity regional AUC data to serve some users who need to prioritize variants with high sensitivity or specificity. (□) To the best of our knowledge, this is the first study to assess the performance of existing methods for non-coding DNMs based on OR values.

Based on the correlation analysis of 24 computational methods, the predictive concordances among the 24 computational methods in the rare somatic variant from COSMIC were obviously higher than in the other three datasets. This may be because somatic variants result from replication errors and DNA damage [50], and hence, somatic variants may have some similar features that germline variants do not, but most variants in the other three datasets are germline variants. Additionally, regBase_CAN [31] in common regulatory variant and disease associated common variant prediction showed a negative correlation with many methods, and most of these methods with a negative correlation with regBase_CAN were those incorporated into regBase_CAN. Compared to other methods, regBase included three methods designed for different purposes and regBase_CAN is one of methods designed for predicting the effects of somatic variants based on somatic variant training dataset [31]. Thus, parameters in regBase_CAN may lead to inconsistent prediction results for common variants with other methods.

Based on our results, we clustered the 24 methods into three groups based on their computational models such as supervised, unsupervised, and semi-supervised models and preliminarily found that ncER (supervised model), DVAR (unsupervised model) and LINSIGH (semi-supervised model) are the representative methods of the above three groups with the highest median of AUC values based on 4 benchmark datasets (**Figure 2**). Additionally, we noted that computational methods showed different prediction efficiencies under different conditions (Figure 2). For example, FATHMM-XF [21] was the best method for the rare germline variant from ClinVar (AUC = 0.8033), but performed poorly for the rare somatic variant from COSMIC (AUC = 0.5933). Although the performance of the computational methods varied for the four different benchmark datasets, the best performance was recorded for the rare germline variant from ClinVar. These results are consistent with those of a previous study [35] and may be attributed to the following reasons. First, most computational methods selected more germline than somatic variants, which may have different genomic features; this selection bias in training data may lead to better performance of the methods in the rare germline variant from ClinVar. Second, it is well-known that genetic variation in many complex quantitative traits results from the joint small effects of multiple variants [51, 52], and non-coding variants often have a weak effect on complex traits [53]. The stronger functional effects of germline variants in the ClinVar [38] database made it easier to distinguish functional variants for these computational methods. Given that the contribution of single expression quantitative trait loci (eQTL) and GWAS SNV to heritability is small, functional prediction of these SNVs remains an enormous challenge.

**Figure 2.**
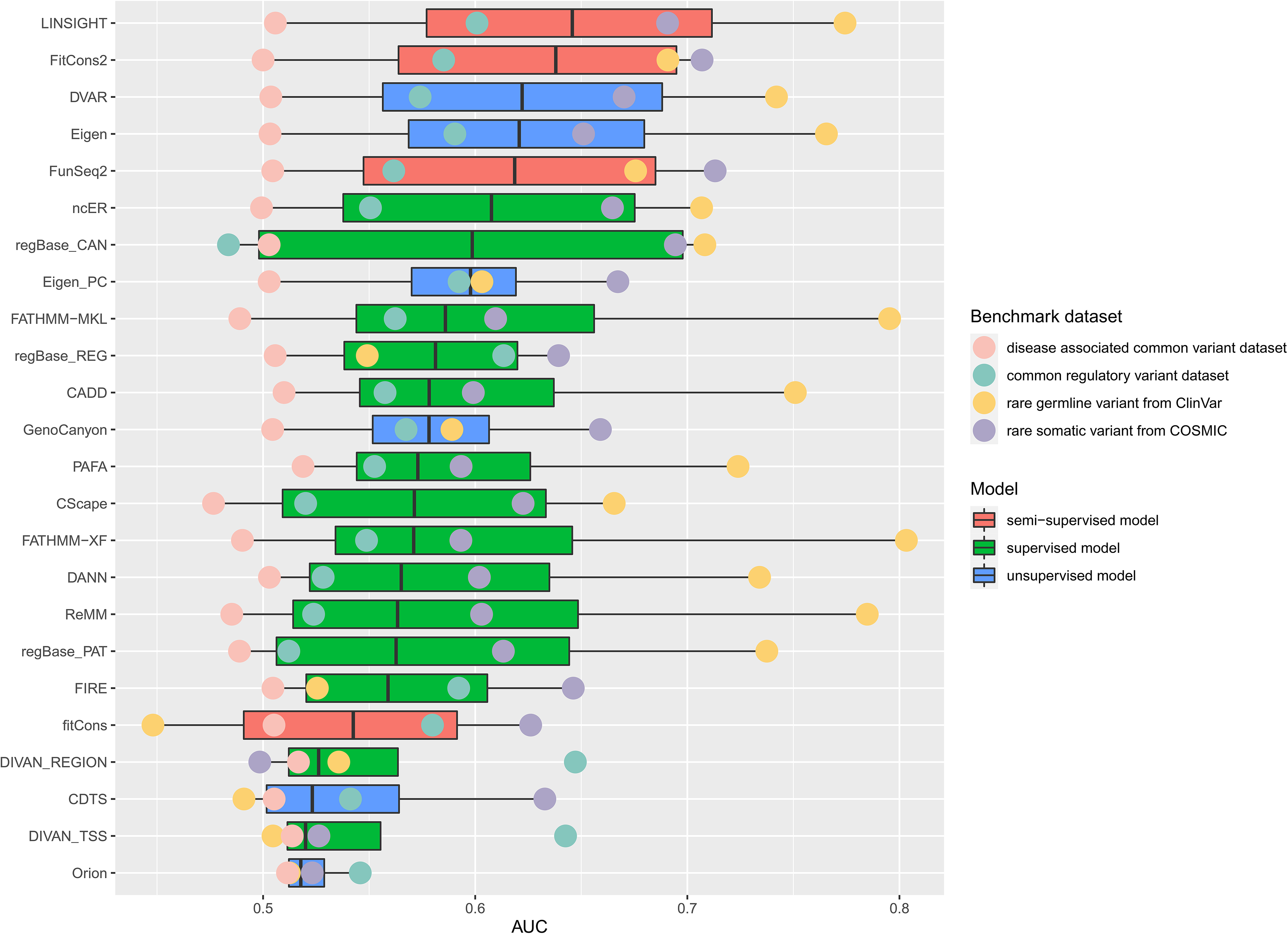
Overall AUCs of four benchmark datasets. AUC, area under the curve. Distributions of AUCs for 24 methods shown in a boxplot. Different coloured balls represent different benchmark datasets. Different coloured bars represent different models

In addition, we found that methods based on supervised models performed better than those based on unsupervised and semi-supervised models in the rare germline variant from ClinVar. This may be explained by the selection of training data, as supervised learning demands representative and correctly labelled training data [54], and many methods based on supervised models select high-quality germline variants sourced data from the HGMD [55] and ClinVar [38] databases as training data. Thus, many methods based on supervised models performed better with the rare germline variant from ClinVar. Furthermore, methods based on semi-supervised models performed better than unsupervised and supervised models in rare somatic variant from COSMIC. This may be because semi-supervised models select both labelled and unlabelled data with stronger and weaker functional effects, respectively, as their training data, whereas the supervised and unsupervised models select labelled and unlabelled data, respectively, as their training data [54].

According to the performance measurement strategy, we divided the 24 methods into three groups (□, □, and □) based on the rank of their AUC values, and every group contained 8 methods (Table S8). None of these methods performed well in all evaluations. This may be because different evaluations represent different aspects of methods performance. Hence, appropriate methods should be selected based on different requirements. In addition, the AUC is not affected by different cut-off values and does not vary significantly with the different ratios of positive and negative variants in benchmark data; thus, we selected the AUC as our major measure.

As we all known, Non-coding DNMs play important roles in neurodevelopmental disorders, such as ASD [46–48], but there is no authoritative database for validated pathogenic DNMs. To assess the prediction performance of the 24 methods for non-coding DNMs, we downloaded non-coding DNMs from patients with ASD and unaffected siblings from our previous study [49]. Although pathogenicity of these DNMs is unclear, the number of pathogenic DNMs from patients with ASD should be more than unaffected siblings. Hence, we selected odds ratios and *P* values to assess the performance of these methods. In addition, we tried our best to collect 57 experimentally validated non-coding transcriptional-regulation-disruption DNMs from ASD probands and 50 nearest non-coding non-pathogenic DNMs in the siblings of ASD patients as our testing dataset to further assess the performance of 24 methods for DNMs. We noted that DVAR, regBase_CAN and FitCons2 showed the better performance with AUC value > 0.77 (table S7). Based on these results, we thought it is still a challenge to make a precious prediction for DNMs.

In this study, we noted that three of 24 compared methods were ensemble prediction models and found that the performances of the three methods (regBase_REG, regBase_PAT, and regBase_CAN) were moderate compared to other methods. In addition, we selected the top 10 methods of each benchmark dataset based on sum of sensitivity and specificity to evaluate whether combined prediction would improve performance. If a variant was predicted as positive by more than half of methods, it was considered as a positive variant. Finally, we assessed the performance of this combined prediction based on the accuracy and MCC and found that combined prediction did not further improve performance. This indicated that it is still a challenge to improve prediction performance for non-coding variants based on existing ensemble models. Hence, we think that more attention should be paid to improving the quality of training data and models to get better prediction performance for non-coding variants.

This study had some limitations. First, there was some potential circularity between the testing and training data of the computational prediction methods [56]. To eliminate potential circularity, we selected testing data that were recorded after 2019 and, as much as possible, removed variants that overlapped with publicly available training data when comparing methods. Given that some methods only provide the source and version without including the exact variants of the training data, a small amount of the benchmark data may still be the same as the training data in the methods. Hence, we suggest that scientists who develop new methods should publish their original training and testing data. Second, although the testing data downloaded from the ClinVar [38], COSMIC [39], GTEx Portal [41], and GWAS Catalog [57] databases have been widely used to develop computational methods and assess their performance, relatively little is known about the functional consequences of variation in the non-coding region of the genome, and most variants in benchmark datasets were not experimentally validated; as such, incorrectly labelled data may have been included in our benchmark data. Therefore, we strongly recommend that scientists select experimentally validated or high-confidence training data develop new methods in future studies.

Taken together, our findings suggest that existing computational methods show acceptable performances only for germline variants and that their predictive ability must be improved for different types of non-coding variants. We strongly recommend that more attention should be paid to the quality of learning data in future software development work. For example, methods should use various training data and genomic features to avoid selection bias. Our findings will serve as a useful guide for clinicians and researchers in choosing appropriate methods for non-coding variant prediction and lead to the development of new methods.

## Materials and methods

### Computational methods and prediction score processing

We compared 24 computational methods that provide precomputed prediction scores for the whole human genome, including 14 methods based on supervised models, 6 methods based on unsupervised models and 4 methods based on semi-supervised models (Table 1). The genomic positions of all precomputed scores were based on GRCh37/hg19. For standardisation, all precomputed scores recorded by interval-level values were transformed into base-wise positions, and each base-wise position was assigned the same score. In addition, these raw scores were transformed into PHRED-scaled scores (−10 × log10[rank/total]) according to the genome-wide distribution of scores for approximately 9 × 10^9^ potential SNVs, which is the set of all three non-reference alleles at each position of the reference assembly. PHRED-scaled scores provide a comparable unit and unifying estimation standard for assessment. For instance, if a raw score in the top 10% of all possible reference genome SNVs, it was represented as a PHRED-scaled score of _≥_10, and a raw score in the top 1‰, was represented as a score of _≥_30. We calculated the mean of the precomputed base-level whole genome DIVAN scores across 45 diseases for both region-matched and transcription start site (TSS)-matched criteria, and then transformed them into a PHRED-scaled score. Other raw and PHRED-scaled scores for all methods were downloaded from a previous study [31] except for DIVAN_TSS and DIVAN_REGION.

### Benchmark datasets of non-coding variants

To evaluate the performance of the 24 methods, it was essential to construct an independent test of datasets in which variants overlapping with the training data were removed from the compared methods as much as possible. Four non-coding independent benchmark datasets were used to assess the performance of the 24 computational methods, including (□) rare germline variant from ClinVar, (□) rare somatic variant from COSMIC, (□) common regulatory variant dataset, and (□) disease associated common variant dataset, and both positive and negative non-coding variants were included in each benchmark dataset (Table 2 and Table S1). We adopted the following strategies to reduce overlap between testing benchmark data and training data for further analysis. First, as all training datasets used were published before 2019, we selected variants recorded in public databases [38, 39] after 2019 to reduce overlap. Second, we comprehensively collected public training data on existing methods and removed overlap between benchmark data and available training data of the computational methods.

The first benchmark dataset (rare germline variant from ClinVar) was downloaded from the ClinVar database [38]. According to the American College of Medical Genetics and Genomics guidelines [58], the variants were classified as ‘pathogenic’, ‘likely pathogenic’, ‘benign’, ‘likely benign’, and ‘uncertain significance’ in the ClinVar database. Furthermore, the ClinVar database contains interpretations of allele origins, and records in ClinVar with ORIGIN = 1 indicated that these variants are germline variants. To improve the accuracy of the benchmark dataset and eliminate overlap between testing benchmark data and training data used in the 24 computational methods, we selected all ‘pathogenic’, ‘likely pathogenic’, and ‘benign’ non-coding germline variants deposited in the ClinVar database after January 2, 2019 as positive and negative variants, respectively. Furthermore, we determine the allele frequency (AF) of these variants based on GnomAD database and noticed that (□) over 80% of ‘pathogenic’ and ‘likely pathogenic’ variants were not observed, (□) over 98% of ‘pathogenic’ and ‘likely pathogenic’ variants had AFs < 0.1%, (□) over 99% of ‘benign’ variants were observed, and (□) over 98% of ‘benign variants had AFs _≥_0.1%. Based on the AF of these variants, we regarded all ‘pathogenic’ and ‘likely pathogenic’ variants as rare variants with AFs < 0.1%. Finally, we only selected all ‘pathogenic’, ‘likely pathogenic’, and ‘benign’ variants with AFs < 0.1% (515 and 1850) as our testing data.

The second benchmark dataset (rare somatic variant from COSMIC) was downloaded from the COSMIC database [39]. As most deleterious non-coding somatic variants are unknown and one criterion for identifying cancer driver variants is to examine their mutational recurrence across multiple samples [59], non-coding somatic variants from the COSMIC database after March 19, 2019 were divided into positive and negative variants, respectively, according to the recurrence of the variant. To increase the reliability of these variants, we also ensured that our positive variants are located on risk genes collected from the CNCDatabase [40], whereas negative variants are not. A total of 2346 variants and 648,471 variants were categorised as positive and negative variants when the threshold value of recurrence was equal to 2 and that 84% of positive variants and 92% of negative variants had AFs < 0.1% based on the GnomAD database. It is widely accepted that most somatic variants observed in the cancer genome are rare [60], and thus we only selected variants with AFs < 0.1% (1966 and 597,222) as our final testing data.

It is well known that non-coding variants influence phenotypes mainly through regulating gene expression level. Hence, we select common regulatory variant dataset as our third benchmark dataset (common regulatory variant dataset) to assess 24 methods. Here, we integrated three independent eQTL test datasets from two studies [18, 31] and removed eight variants that were labelled differently in both studies as our testing data. The positive dataset included (□) high-confidence eQTL single-nucleotide polymorphisms from the GTEx Portal database [41] and (□) multi-tissue eQTL single-nucleotide polymorphisms fine-mapping data from Brown’s study [42] and the GTEx Portal database [41]. The negative dataset was randomly sampled by vSampler [61] based on 1000 Genomes Project phase3 (1000G P3) [43] and negative variants had matched minor allele frequencies (MAFs), distance to closest transcription start site, gene density, and number of variants in LD (Table S9). Notably, all positive and negative variants are non-coding, with MAFs > 5% based on 1000G P3. We also referred to the criteria of test sets from Li’s study [36] and only included paired positive and negative variants beyond 1 kbps from each other as our final testing data to prevent physically proximate variants from confounding.

The fourth benchmark dataset (disease associated common variant dataset) was downloaded from the CAUSALdb database [44] and 1000 Genomes Project [43]. We only selected non-coding SNVs in the intersection set of credible sets defined by three tools including PAINTOR [62], CAVIARBF [63], and FINEMAP [64] with MAFs > 5% based on the 1000 Genomes Project as positive variants and corresponding non-coding SNVs in the same LD blocks with r^2^ thresholds > 0.2 from the 1000 Genomes Project with MAFs > 5% as negative variants. Overlapping variants between positive and negative data as well as positive variants without corresponding negative variants were excluded from the analysis.

### Correlation analysis

Spearman rank correlation coefficient was used to evaluate the relationships among the 24 compared computational methods based on the 4 non-coding benchmark datasets described above. Specifically, Spearman rank correlation coefficients were calculated between any two computational methods for each benchmark dataset, in which variants with missing values for a method were excluded and the results of correlation analyses were visualised in the form of heat maps. In addition, for each benchmark dataset, we performed correlation analysis based on the positive and negative variant datasets.

### Measures for performance evaluation

The performances of the 24 computational methods were assessed based on the following 12 criteria: (□) the positive predictive value, as the proportion of positive results in the computational methods that was positive under the benchmark dataset; (□) the negative predictive value, the proportion of negative results in computational methods that was negative under the benchmark dataset; (□) the false-negative rate, which is calculated as the ratio of the number of positive events wrongly categorised as negative by the computational method to the total number of actual positive events under the benchmark dataset; (□) the sensitivity (or true-positive rate), which measures the proportion of actual positives under the benchmark dataset that are correctly identified as such by the computational method. The false-negative rate and sensitivity are paired measures with a sum equal to 100%; (□) the false-positive rate, calculated as the ratio of the number of negative events wrongly categorised as positive by the computational method to the total number of actual negative events under the benchmark dataset; (□) the specificity (or true-negative rate), which measures the proportion of actual negatives under the benchmark dataset that is correctly identified as such by the computational method. The false-positive rate and specificity are paired measures with a sum equal to 100%; (□) the accuracy, which represents the proportion of positive and negative variants in the benchmark data that are correctly predicted as positive and negative variants, respectively; (□) the Mathews correlation coefficient, as a correlation coefficient (range of −1 to 1) between the observed and predicted classifications, where 1 indicates a perfect prediction, 0 no better than random prediction, and −1 complete disagreement between the prediction and true classification; (□) the ROC curve, a graphical plot that illustrates the predictive ability of a computational method as its discrimination thresholds are varied; (□) the AUC value, which ranges from 0 to 1 for each ROC curve, such that a higher AUC indicates better performance of the computational method; (□) the hser-AUC, which is the AUC corresponding to high sensitivity (true-positive rate > 95%); and (□) the hspr-AUC, which is the AUC corresponding to high specificity (true-negative rate > 95%). The hser-AUC and hspr-AUC were evaluated to serve some users who require a distinction between positive variants with high sensitivity or specificity. Given that many computational methods do not offer recommended cut-off values, all measures described above were calculated based on the best thresholds corresponding to the best sum of sensitivity and specificity. In addition, the best thresholds, sensitivities, specificities, AUCs, hspr-AUCs, and hser-AUCs were calculated using the ‘pROC’ package [65] based on PHRED-scaled scores.

### Non-coding DNMs from the Simons simplex collection (SSC)

Non-coding DNMs identified in 1902 patients with ASD and 1902 unaffected siblings were downloaded from the SSC [47, 66] (Table S1) and were previously catalogued in the Gene4Denovo database that we developed [49]. Comparison of the performance of computational methods for non-coding DNMs was based on PHRED-scaled scores. We compared the burden of functional non-coding variants that were predicted by the computational methods in the ASD and sibling groups under different cut-off values. To assess the performance of computational methods for DNMs, we calculated the ORs, 95% confidence interval of the OR, and *P* value between ASD and unaffected siblings using the two-sided Poisson’s ratio test.

### Experimentally validated non-coding DNMs from ASD

We collected experimentally validated non-coding transcriptional-regulation-disruption DNMs from ASD probands[48] and nearest non-coding non-pathogenic DNMs in the siblings of ASD patients[31] as our supplementary test dataset (Table S1) to further assess the performance of 24 methods for DNMs.

### Code availability

Source code for all analysis and benchmark is available on XXXXX at XXXXXX

## CRediT author statement

Zheng Wang: Conceptualization, Methodology, Validation, Formal analysis, Investigation, Data Curation, Writing - original draft, Writing - review & editing. Guihu Zhao: Conceptualization, Methodology, Software, Data Curation, Writing - original draft. Bin Li: Investigation. Zhenghuan Fang: Methodology. Qian Chen: Investigation. Xiaomeng Wang: Data Curation. Tengfei Luo: Investigation. Yijing Wang: Investigation. Qiao Zhou: Investigation, Data Curation. Kuokuo Li: Visualization. Lu Xia: Investigation. Yi Zhang: Investigation. Xun Zhou: Investigation, Data Curation, Visualization. Hongxu Pan: Investigation, Data Curation, Visualization. Yuwen Zhao: Investigation, Data Curation, Visualization. Yige Wang: Investigation, Data Curation, Visualization. Lin wang: Data Curation. Jifeng Guo: Resources, Supervision, Project administration. Beisha Tang: Conceptualization, Resources, Writing - review & editing Kun Xia: Resources, Supervision, Project administration. Jinchen Li: Conceptualization, Resources, Writing - review & editing, Supervision, Project administration, Funding acquisition.

## Acknowledgements

This work was supported by funds from the National Natural Science Foundation of China (grant number 81801133 to JCL), the Young Elite Scientist Sponsorship Program by CAST (grant number 2018QNRC001 to JCL), the Innovation-Driven Project of Central South University (grant number 20180033040004 to JCL), the Natural Science Foundation for Young Scientists of Hunan Province, China (grant number 2019JJ50974 to GHZ), and the Natural Science Foundation of Hunan Province for outstanding Young Scholars (grant number 2020JJ3059 to JCL).

## Supplementary material

**Figure S1.**
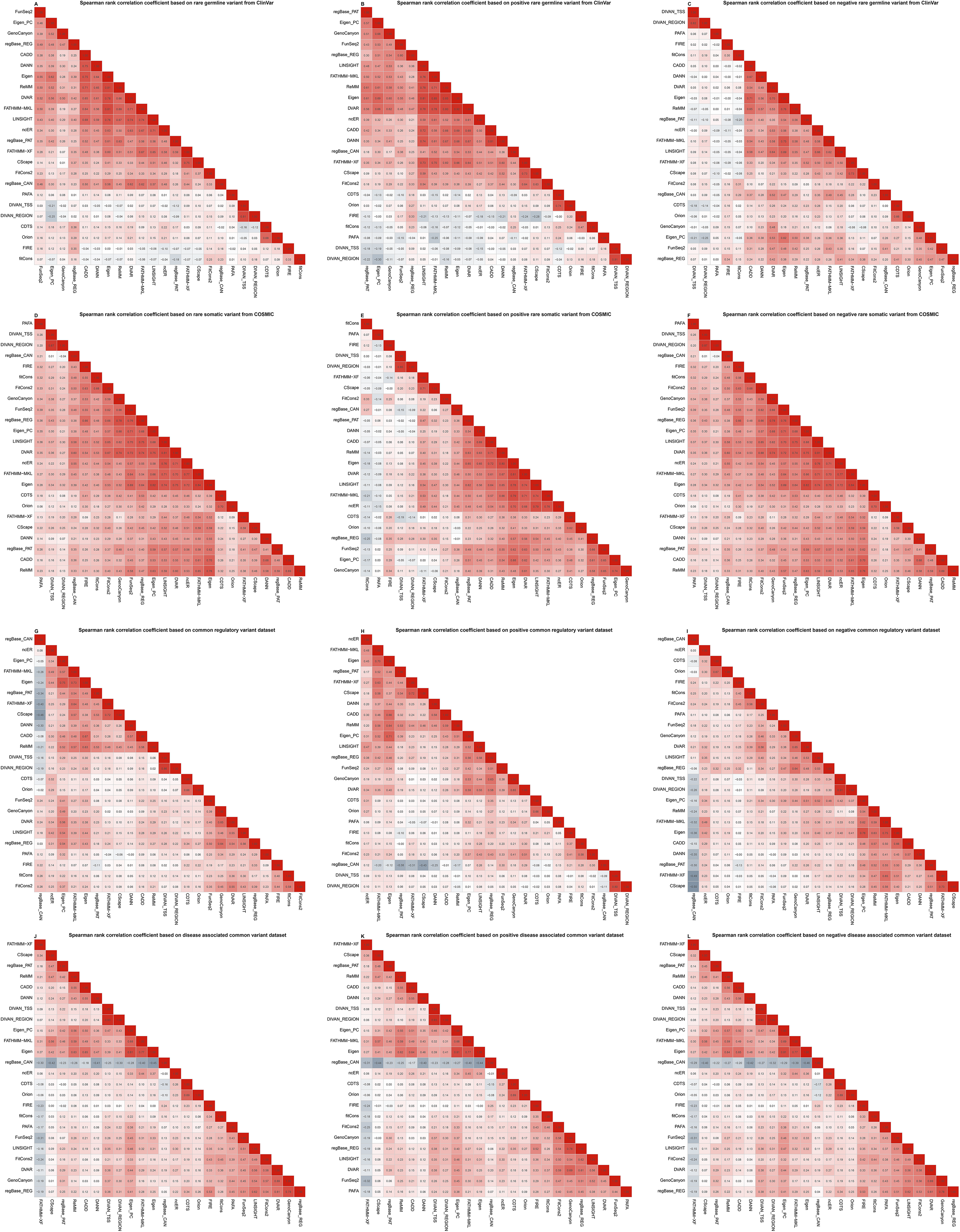
Spearman rank correlation coefficients among 24 computational methods. Spearman rank correlation coefficients were calculated between any two computational methods for four benchmark datasets including their positive variant datasets and negative variant datasets.

**Figure S2.**
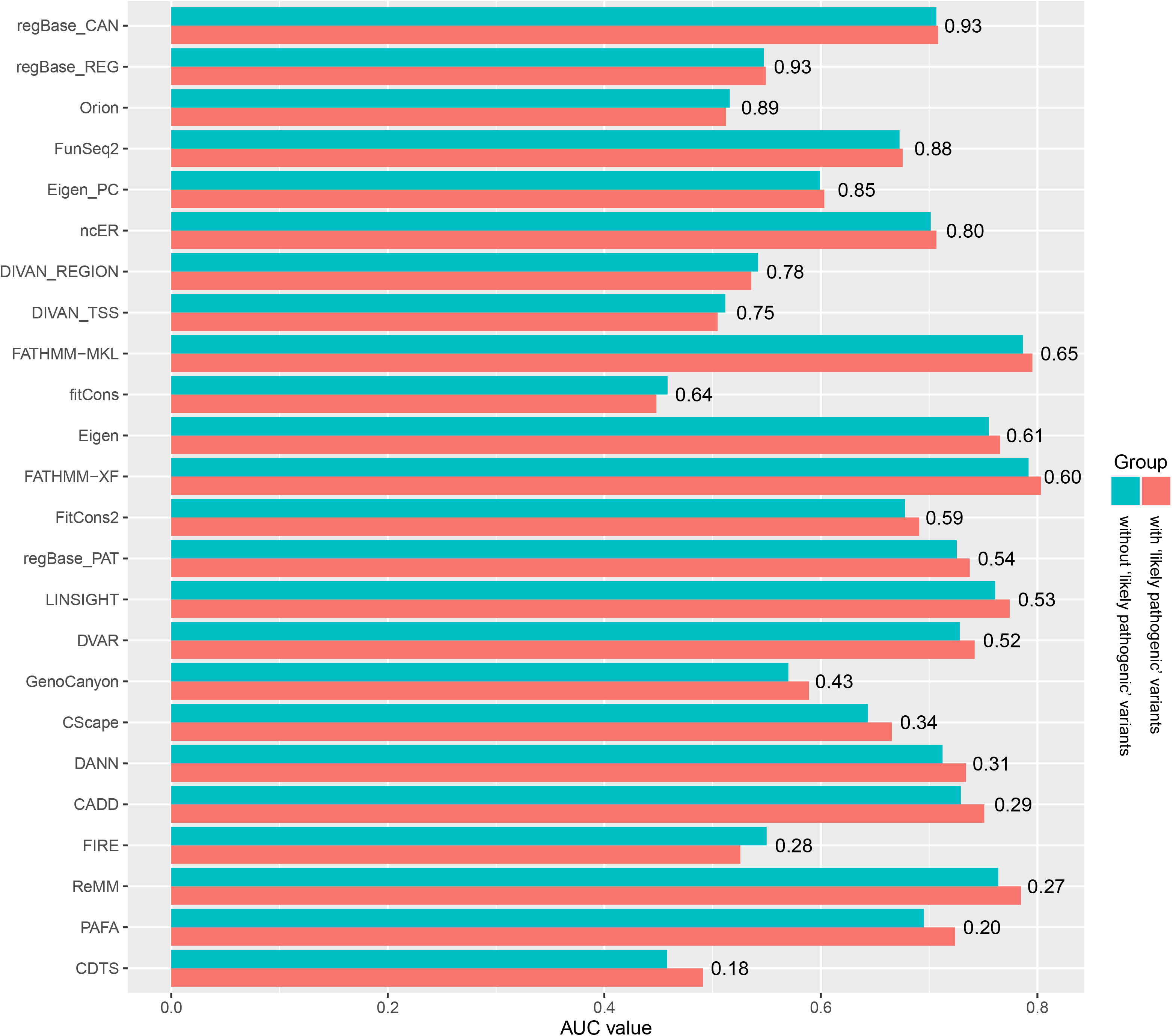
AUCs of 24 computational methods based on different rare germline variant from ClinVar. AUC, area under the curve. Different coloured bars represent methods based on different rare germline variant from ClinVar.

**Figure S3.**
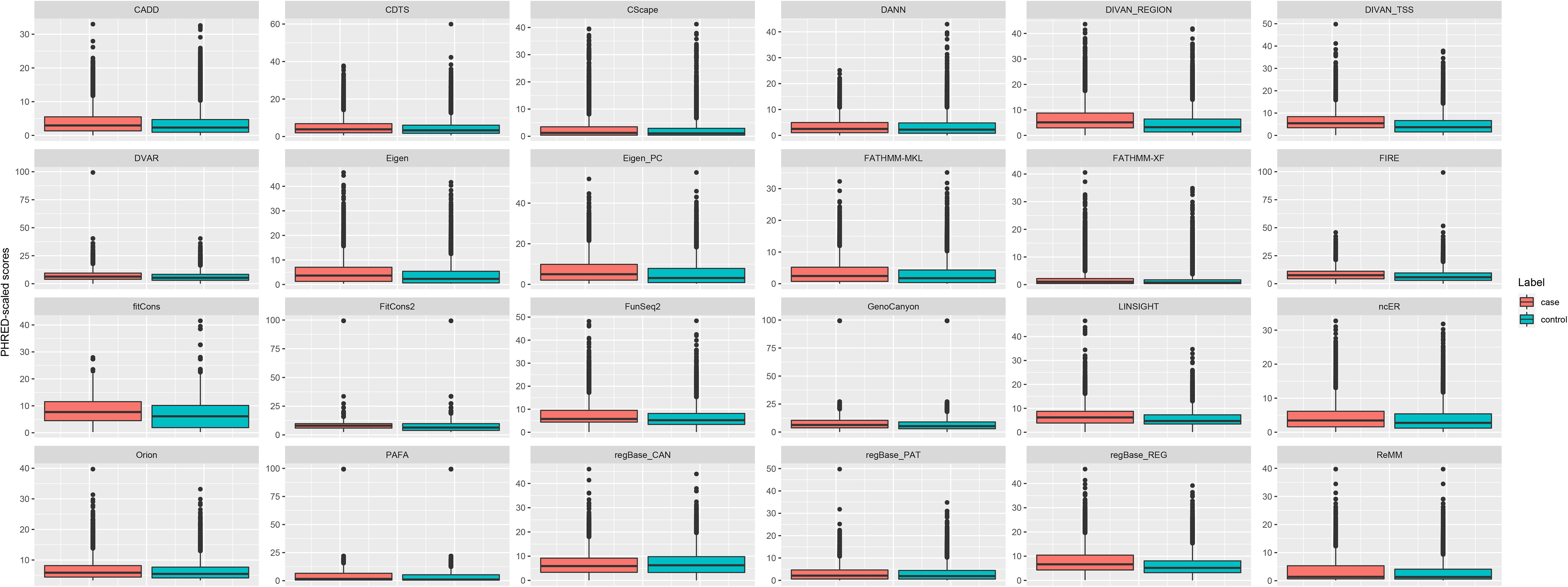
Overall PHRED-scaled scores of 24 computational methods based on the common regulatory variant dataset. Distributions of PHRED-scaled scores for 24 methods shown in a boxplot. Black dots represent outlier values of PHRED-scaled scores.

**Figure S4.**
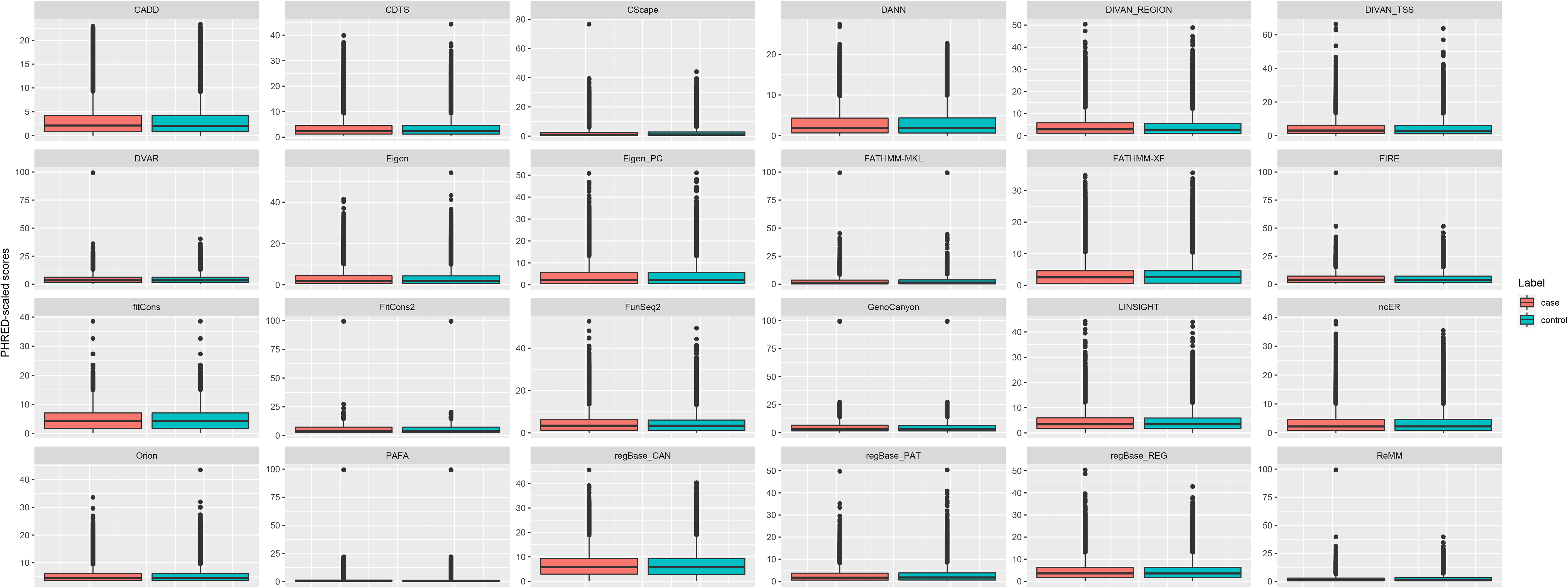
Overall PHRED-scaled scores of 24 computational methods based on the disease associated common variant dataset. Distributions of PHRED-scaled scores for 24 methods shown in a boxplot. Black dots represent outlier values of PHRED-scaled scores.

**Figure S5.**
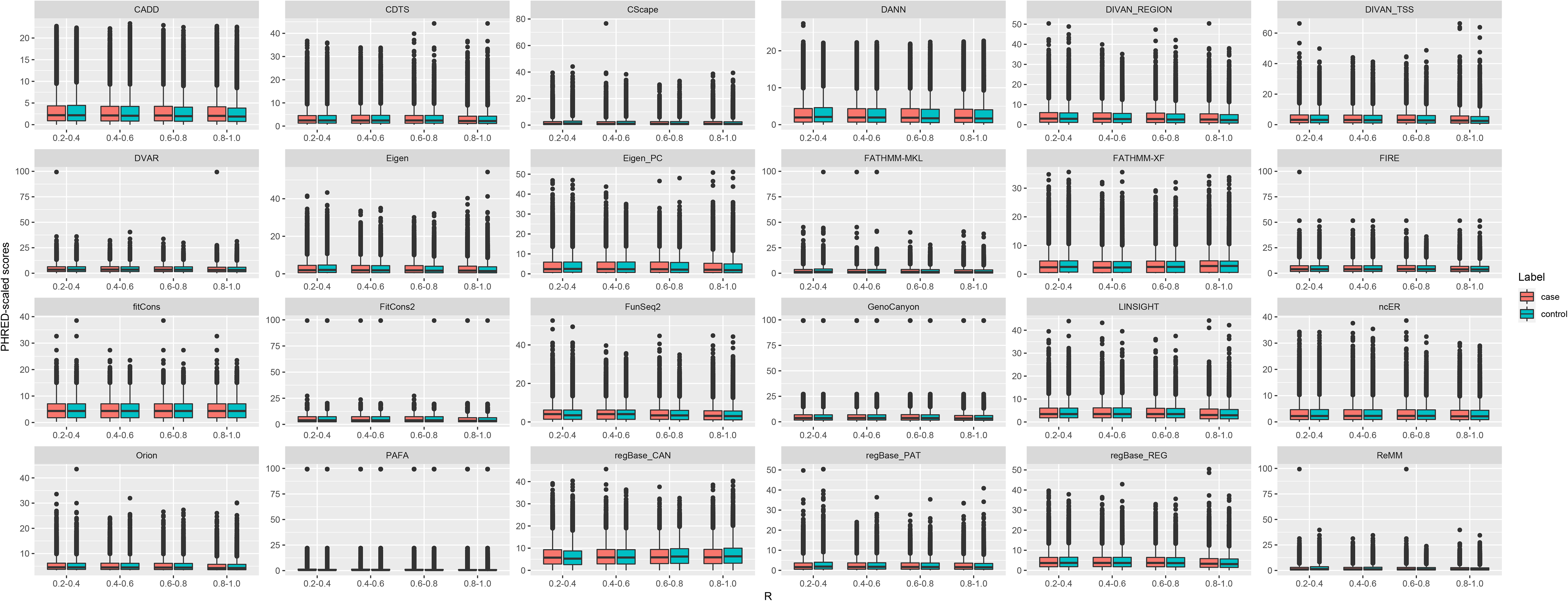
Overall PHRED-scaled scores of 24 computational methods based on different subsets of the disease associated common variant dataset. Distributions of PHRED-scaled scores for 24 methods shown in a boxplot. Black dots represent outlier values of PHRED-scaled scores.

**Figure S6.**
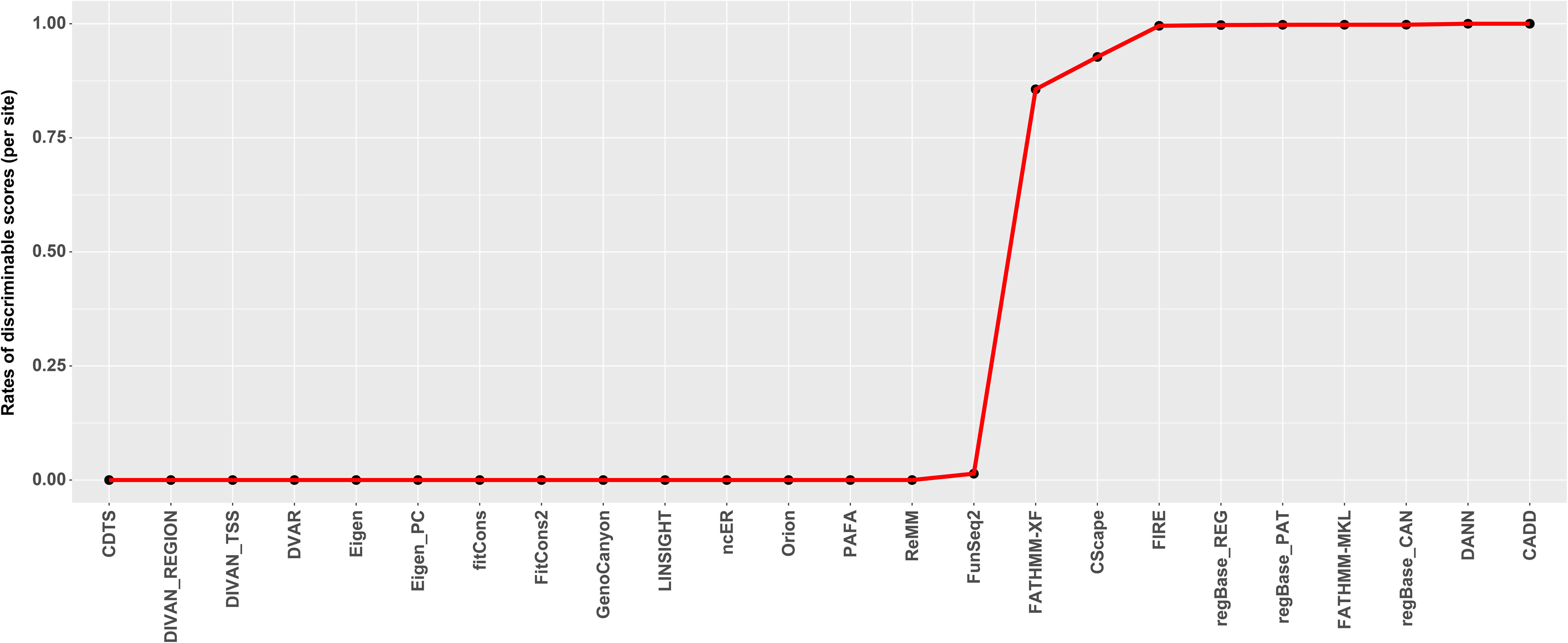
Proportion of discriminable scores (per site) among 24 methods for whole genome.

**Figure S7.**
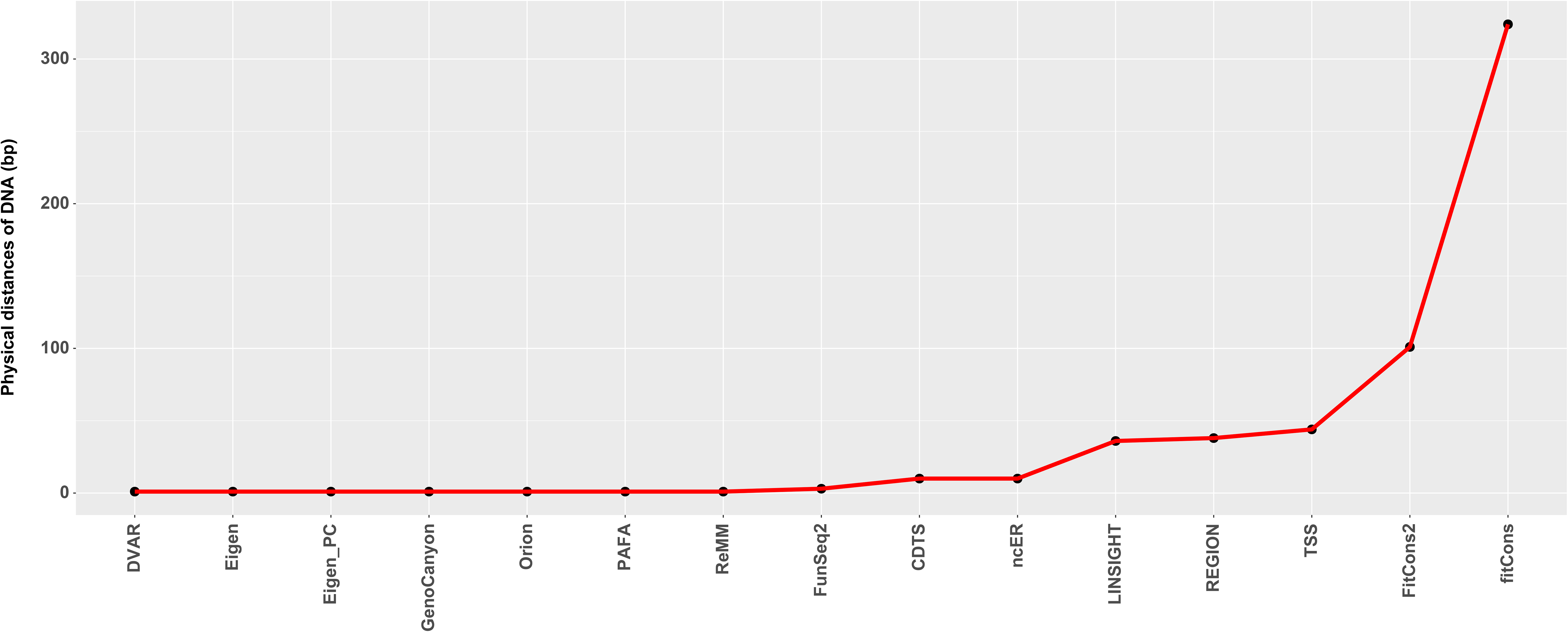
Resolution of prediction scores among 24 methods. We used the cumulative sum of proportions of different physical distances from 1 to largest until it was not smaller than 0.9 then selected the last physical distances as the resolution.

**Figure S8.** Overall AUCs of four benchmark datasets. AUC, area under the curve. Distributions of AUCs for 24 methods shown in a boxplot. Different coloured balls represent different benchmark datasets.

**Table S1.** Testing variants of five independent datasets.

**Table S2.**
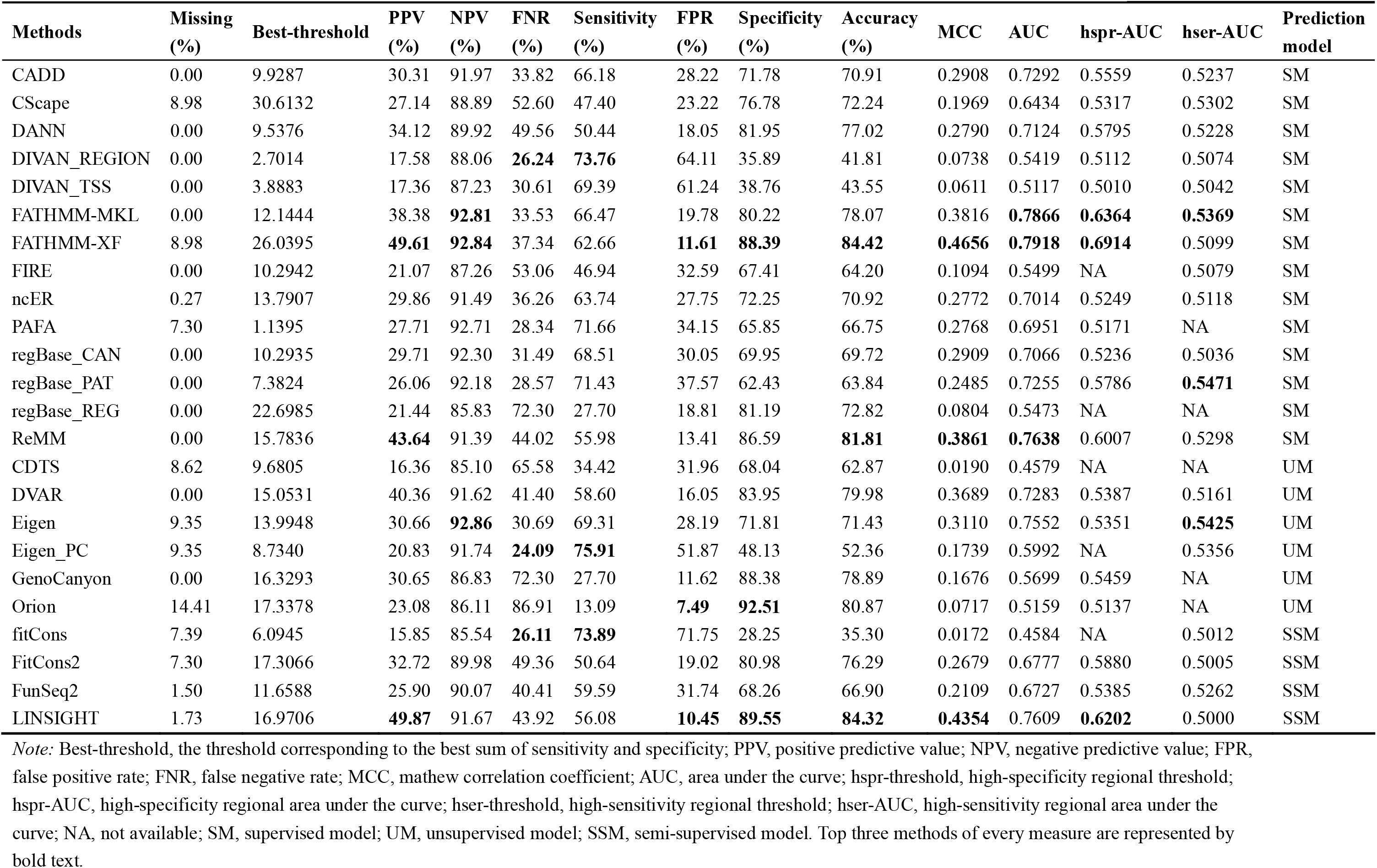
Performance evaluation based on the rare germline variant from ClinVar without ‘likely pathogenic’ variants.

**Table S3.** Performance evaluation based on the rare somatic variant from COSMIC.

| Methods | Missing (%) | Best-threshold | PPV (%) | NPV (%) | FNR (%) | Sensitivity (%) | FPR (%) | Specificity (%) | Accuracy (%) | MCC | AUC | hspr-AUC | hser-AUC | Predict model |
| --- | --- | --- | --- | --- | --- | --- | --- | --- | --- | --- | --- | --- | --- | --- |
| CADD | 0.00 | 6.7405 | 0.49 | 99.76 | 46.64 | 53.36 | 35.39 | 64.61 | 64.58 | 0.0215 | 0.5991 | 0.5082 | NA | SM |
| CScape | 7.32 | 9.1969 | 0.48 | 99.78 | 49.26 | 50.74 | **32.40** | **67.60** | **67.55** | 0.0216 | 0.6226 | 0.5262 | 0.5151 | SM |
| DANN | 0.00 | 7.6538 | 0.52 | 99.74 | 58.60 | 41.40 | **26.03** | **73.97** | **73.87** | 0.0200 | 0.6020 | 0.5289 | 0.5056 | SM |
| DIVAN_REGION | 0.00 | 1.5314 | 0.35 | 99.78 | **12.92** | **87.08** | 80.54 | 19.46 | 19.68 | 0.0094 | 0.4984 | NA | 0.5066 | SM |
| DIVAN_TSS | 0.00 | 2.0320 | 0.38 | 99.83 | **12.46** | **87.54** | 76.31 | 23.69 | 23.90 | 0.0151 | 0.5263 | NA | 0.5087 | SM |
| FATHMM-MKL | 0.02 | 7.6906 | 0.49 | 99.76 | 46.80 | 53.20 | 35.57 | 64.43 | 64.39 | 0.0211 | 0.6096 | 0.5185 | 0.5013 | SM |
| FATHMM-XF | 7.32 | 10.7800 | 0.48 | 99.79 | 44.85 | 55.15 | 34.97 | 65.03 | 65.00 | 0.0234 | 0.5933 | 0.5196 | NA | SM |
| FIRE | 0.00 | 4.9090 | 0.46 | **99.87** | 15.51 | 84.49 | 60.16 | 39.84 | 39.99 | 0.0284 | 0.6462 | 0.5072 | 0.5897 | SM |
| ncER | 0.78 | 6.8488 | 0.51 | 99.83 | 28.30 | 71.70 | 45.83 | 54.17 | 54.22 | 0.0297 | 0.6647 | 0.5135 | 0.5448 | SM |
| PAFA | 2.82 | 6.4113 | 0.48 | 99.78 | 37.26 | 62.74 | 43.53 | 56.47 | 56.49 | 0.0223 | 0.5933 | 0.5196 | NA | SM |
| regBase_CAN | 0.00 | 7.3988 | **0.58** | 99.84 | 30.01 | 69.99 | 39.69 | 60.31 | 60.34 | **0.0354** | **0.6944** | 0.5132 | 0.5707 | SM |
| regBase_PAT | 0.00 | 6.0064 | 0.49 | 99.76 | 47.76 | 52.24 | 35.14 | 64.86 | 64.82 | 0.0205 | 0.6133 | 0.5225 | 0.5033 | SM |
| regBase_REG | 0.00 | 9.2831 | 0.48 | 99.78 | 39.57 | 60.43 | 40.82 | 59.18 | 59.19 | 0.0228 | 0.6393 | 0.5157 | 0.5389 | SM |
| ReMM | 0.00 | 8.0817 | 0.49 | 99.76 | 49.24 | 50.76 | 33.96 | 66.04 | 65.99 | 0.0203 | 0.6030 | 0.5192 | 0.5182 | SM |
| CDTS | 14.24 | 7.7832 | 0.46 | 99.81 | 44.52 | 55.48 | 34.06 | 65.94 | 65.91 | 0.0241 | 0.6328 | 0.5115 | 0.5084 | UM |
| DVAR | 0.00 | 6.9717 | 0.51 | 99.84 | 25.38 | 74.62 | 47.76 | 52.24 | 52.31 | 0.0307 | 0.6702 | 0.5111 | 0.5645 | UM |
| Eigen | 5.96 | 9.0256 | 0.50 | 99.82 | 37.62 | 62.38 | 37.69 | 62.31 | 62.31 | 0.0281 | 0.6511 | 0.5232 | 0.5060 | UM |
| Eigen_PC | 5.96 | 7.2917 | 0.49 | 99.84 | 28.84 | 71.16 | 44.63 | 55.37 | 55.42 | 0.0294 | 0.6673 | 0.5243 | 0.5133 | UM |
| GenoCanyon | 0.00 | 7.8766 | **0.56** | 99.78 | 46.29 | 53.71 | **31.57** | **68.43** | **68.39** | 0.0272 | 0.6590 | **0.5368** | 0.5374 | UM |
| Orion | 27.71 | 7.9485 | 0.46 | 99.65 | 46.08 | 53.92 | 46.82 | 53.18 | 53.18 | 0.0090 | 0.5232 | 0.5001 | NA | UM |
| fitCons | 2.93 | 12.0551 | 0.55 | 99.79 | 41.70 | 58.30 | 35.27 | 64.73 | 64.71 | 0.0277 | 0.6262 | **0.5331** | NA | SSM |
| FitCons2 | 2.82 | 11.3451 | **0.59** | 99.84 | 28.46 | 71.54 | 40.55 | 59.45 | 59.49 | **0.0364** | **0.7069** | 0.5325 | **0.5942** | SSM |
| FunSeq2 | 3.97 | 4.5040 | 0.47 | **99.90** | 17.02 | 82.98 | 51.70 | 48.30 | 48.41 | **0.0340** | **0.7131** | **0.5487** | **0.6205** | SSM |
| LINSIGHT | 4.69 | 6.1970 | 0.46 | **99.90** | **14.83** | **85.17** | 55.69 | 44.31 | 44.43 | 0.0325 | 0.6907 | 0.5163 | **0.5967** | SSM |

*Note:* Best-threshold, the threshold corresponding to the best sum of sensitivity and specificity; PPV, positive predictive value; NPV, negative predictive value; FPR, false positive rate; FNR, false negative rate; MCC, mathew correlation coefficient; AUC, area under the curve; hspr-threshold, high-specificity regional threshold; hspr-AUC, high-specificity regional area under the curve; hser-threshold, high-sensitivity regional threshold; hser-AUC, high-sensitivity regional area under the curve; NA, not available; SM, supervised model; UM, unsupervised model; SSM, semi-supervised model. Top three methods of every measure are represented by bold text.

**Table S4.** Performance evaluation based on the common regulatory variant dataset.

| Methods | Missing (%) | Best-threshold | PPV (%) | NPV (%) | FNR (%) | Sensitivity (%) | FPR (%) | Specificity (%) | Accuracy (%) | MCC | AUC | hspr-AUC | hser-AUC | Predict model |
| --- | --- | --- | --- | --- | --- | --- | --- | --- | --- | --- | --- | --- | --- | --- |
| CADD | 0.00 | 1.8401 | 53.57 | 55.84 | 33.52 | 66.48 | 57.61 | 42.39 | 54.43 | 0.0914 | 0.5575 | 0.5021 | 0.5102 | SM |
| CScape | 1.51 | 3.1122 | 55.12 | 50.85 | 72.11 | 27.89 | **23.33** | **76.67** | 51.95 | 0.0522 | 0.5201 | 0.5058 | 0.5022 | SM |
| DANN | 0.00 | 0.7594 | 51.72 | 56.43 | 18.38 | 81.62 | 76.19 | 23.81 | 52.71 | 0.0665 | 0.5283 | NA | 0.5142 | SM |
| DIVAN_REGION | 0.00 | 2.4827 | **58.20** | **69.58** | **17.95** | **82.05** | 58.93 | 41.07 | **61.56** | **0.2534** | **0.6472** | **0.5104** | **0.5723** | SM |
| DIVAN_TSS | 0.00 | 2.9377 | 58.15 | **69.30** | **18.24** | **81.76** | 58.84 | 41.16 | **61.46** | **0.2508** | **0.6426** | 0.5066 | **0.5714** | SM |
| FATHMM-MKL | 0.00 | 1.5323 | 54.30 | 55.93 | 37.06 | 62.94 | 52.96 | 47.04 | 54.99 | 0.1010 | 0.5623 | 0.5004 | 0.5152 | SM |
| FATHMM-XF | 1.51 | 1.5933 | 57.30 | 52.41 | 64.04 | 35.96 | **27.54** | **72.46** | 53.97 | 0.0905 | 0.5488 | **0.5138** | 0.5003 | SM |
| FIRE | 0.00 | 5.8832 | 56.37 | 58.16 | 36.70 | 63.30 | 48.99 | 51.01 | 57.15 | 0.1442 | 0.5922 | 0.5074 | 0.5179 | SM |
| ncER | 0.34 | 2.7862 | 54.02 | 54.63 | 41.93 | 58.07 | 49.47 | 50.53 | 54.30 | 0.0862 | 0.5506 | 0.5001 | 0.5088 | SM |
| PAFA | 0.00 | 1.6172 | 54.67 | 53.89 | 50.32 | 49.68 | 41.19 | 58.81 | 54.25 | 0.0853 | 0.5526 | 0.5054 | NA | SM |
| regBase_CAN | 0.00 | 1.6270 | 50.22 | 52.02 | **9.50** | **90.50** | 89.70 | 10.30 | 50.40 | 0.0134 | 0.4837 | NA | 0.5022 | SM |
| regBase_PAT | 0.00 | 2.9755 | 52.06 | 51.30 | 59.72 | 40.28 | 37.10 | 62.90 | 51.59 | 0.0327 | 0.5121 | NA | NA | SM |
| regBase_REG | 0.00 | 5.7329 | 57.49 | 57.87 | 41.10 | 58.90 | 43.55 | 56.45 | 57.67 | 0.1535 | **0.6134** | **0.5148** | **0.5262** | SM |
| ReMM | 0.00 | 5.4273 | 55.96 | 51.69 | 75.29 | 24.71 | **19.44** | **80.56** | 52.63 | 0.0635 | 0.5238 | 0.5057 | 0.5048 | SM |
| CDTS | 9.68 | 3.4039 | 55.45 | 51.14 | 44.83 | 55.17 | 48.57 | 51.43 | 53.38 | 0.0660 | 0.5412 | 0.5031 | 0.5059 | UM |
| DVAR | 0.00 | 4.6740 | 54.73 | 57.70 | 32.17 | 67.83 | 56.12 | 43.88 | 55.86 | 0.1207 | 0.5740 | 0.5023 | 0.5116 | UM |
| Eigen | 0.75 | 2.7022 | 57.24 | 57.42 | 39.86 | 60.14 | 45.53 | 54.47 | 57.33 | 0.1464 | 0.5904 | 0.5072 | 0.5143 | UM |
| Eigen_PC | 0.75 | 2.9487 | 56.83 | **59.09** | 33.15 | 66.85 | 51.46 | 48.54 | 57.75 | 0.1565 | 0.5924 | 0.5064 | 0.5173 | UM |
| GenoCanyon | 0.00 | 5.2397 | 54.40 | 55.35 | 40.29 | 59.71 | 50.06 | 49.94 | 54.83 | 0.0970 | 0.5674 | 0.5073 | 0.5160 | UM |
| Orion | 38.34 | 4.6326 | **60.30** | 47.42 | 29.32 | 70.68 | 63.76 | 36.24 | 56.15 | 0.0731 | 0.5458 | 0.5030 | 0.5124 | UM |
| fitCons | 0.02 | 8.6616 | **58.55** | 54.98 | 56.90 | 43.10 | 30.51 | 69.49 | 56.30 | 0.1305 | 0.5799 | NA | 0.5010 | SSM |
| FitCons2 | 0.00 | 6.8789 | 56.44 | 58.01 | 37.43 | 62.57 | 48.28 | 51.72 | 57.14 | 0.1437 | 0.5851 | NA | 0.5128 | SSM |
| FunSeq2 | 1.43 | 5.2371 | 54.77 | 54.05 | 41.95 | 58.05 | 49.28 | 50.72 | 54.44 | 0.0880 | 0.5616 | 0.5085 | 0.5056 | SSM |
| LINSIGHT | 1.51 | 4.7214 | 57.41 | 58.44 | 34.80 | 65.20 | 49.71 | 50.29 | **57.85** | **0.1567** | 0.6009 | 0.5089 | 0.5161 | SSM |

*Note:* Best-threshold, the threshold corresponding to the best sum of sensitivity and specificity; PPV, positive predictive value; NPV, negative predictive value; FPR, false positive rate; FNR, false negative rate; MCC, mathew correlation coefficient; AUC, area under the curve; hspr-threshold, high-specificity regional threshold; hspr-AUC, high-specificity regional area under the curve; hser-threshold, high-sensitivity regional threshold; hser-AUC, high-sensitivity regional area under the curve; NA, not available; SM, supervised model; UM, unsupervised model; SSM, semi-supervised model. Top three methods of every measure are represented by bold text.

**Table S5.** Performance evaluation based on the disease associated common variant dataset.

| Methods | Missing (%) | Best-threshold | PPV (%) | NPV (%) | FNR (%) | Sensitivity (%) | FPR (%) | Specificity (%) | Accuracy (%) | MCC | AUC | hspr-AUC | hser-AUC | Prediction model |
| --- | --- | --- | --- | --- | --- | --- | --- | --- | --- | --- | --- | --- | --- | --- |
| CADD | 0.00 | 1.3843 | 0.50 | 0.52 | 0.36 | 0.64 | 0.63 | 0.37 | **0.51** | 0.0182 | 0.5099 | 0.5002 | 0.5020 | SM |
| CScape | 0.00 | 27.2206 | **0.59** | 0.51 | 1.00 | 0.00 | **0.00** | **1.00** | **0.51** | 0.0059 | 0.4766 | NA | NA | SM |
| DANN | 0.00 | 0.2798 | 0.50 | **0.53** | **0.13** | **0.87** | 0.86 | 0.14 | 0.50 | 0.0187 | 0.5030 | NA | 0.5028 | SM |
| DIVAN_REGION | 0.00 | 2.0742 | 0.50 | 0.52 | 0.39 | 0.61 | 0.58 | 0.42 | **0.51** | **0.0259** | **0.5167** | **0.5014** | **0.5036** | SM |
| DIVAN_TSS | 0.00 | 1.8537 | 0.50 | 0.52 | 0.35 | 0.65 | 0.63 | 0.37 | **0.51** | 0.0216 | **0.5137** | 0.5007 | **0.5037** | SM |
| FATHMM-MKL | 0.00 | 10.4953 | 0.50 | 0.51 | 0.95 | 0.05 | 0.05 | 0.95 | **0.51** | 0.0043 | 0.4890 | 0.5005 | NA | SM |
| FATHMM-XF | 0.00 | 4.8704 | 0.49 | 0.51 | 0.77 | 0.23 | 0.23 | 0.77 | **0.51** | 0.0017 | 0.4903 | 0.5001 | NA | SM |
| FIRE | 0.00 | 8.0711 | 0.50 | 0.51 | 0.78 | 0.22 | 0.21 | 0.79 | **0.51** | 0.0104 | 0.5046 | 0.5009 | 0.5002 | SM |
| ncER | 0.00 | 11.3622 | **0.51** | 0.51 | 0.96 | 0.04 | 0.04 | 0.96 | **0.51** | 0.0091 | 0.4993 | **0.5010** | 0.5003 | SM |
| PAFA | 0.00 | 0.5155 | **0.51** | **0.53** | 0.44 | 0.56 | 0.52 | 0.48 | **0.52** | **0.0332** | **0.5188** | 0.5002 | 0.5023 | SM |
| regBase_CAN | 0.00 | 2.5935 | 0.49 | 0.52 | 0.22 | 0.78 | 0.77 | 0.23 | 0.50 | 0.0087 | 0.5029 | NA | 0.5007 | SM |
| regBase_PAT | 0.00 | 14.6167 | 0.50 | 0.51 | 0.99 | 0.01 | **0.01** | **0.99** | **0.51** | 0.0007 | 0.4889 | NA | NA | SM |
| regBase_REG | 0.00 | 3.0243 | 0.50 | 0.51 | 0.43 | 0.57 | 0.56 | 0.44 | 0.50 | 0.0113 | 0.5057 | **0.5010** | 0.5005 | SM |
| ReMM | 0.00 | 37.0900 | **0.75** | 0.51 | 1.00 | 0.00 | **0.00** | **1.00** | **0.51** | 0.0027 | 0.4853 | NA | NA | SM |
| CDTS | 0.17 | 0.8700 | 0.50 | 0.52 | **0.12** | **0.88** | 0.87 | 0.13 | 0.50 | 0.0147 | 0.5052 | 0.5003 | 0.5019 | UM |
| DVAR | 0.00 | 2.9796 | 0.50 | 0.51 | 0.45 | 0.55 | 0.55 | 0.45 | 0.50 | 0.0086 | 0.5036 | 0.5008 | NA | UM |
| Eigen | 0.00 | 2.7579 | 0.50 | 0.51 | 0.60 | 0.40 | 0.39 | 0.61 | **0.51** | 0.0115 | 0.5033 | 0.5009 | NA | UM |
| Eigen_PC | 0.00 | 2.3400 | 0.50 | 0.51 | 0.50 | 0.50 | 0.49 | 0.51 | 0.50 | 0.0076 | 0.5029 | 0.5004 | 0.5001 | UM |
| GenoCanyon | 0.00 | 2.6909 | 0.50 | 0.51 | 0.40 | 0.60 | 0.60 | 0.40 | 0.50 | 0.0086 | 0.5045 | 0.5002 | 0.5006 | UM |
| Orion | 0.55 | 3.3963 | 0.51 | **0.56** | **0.06** | **0.94** | 0.92 | 0.08 | **0.51** | **0.0358** | 0.5115 | 0.5007 | **0.5063** | UM |
| fitCons | 0.00 | 4.5007 | 0.50 | 0.51 | 0.58 | 0.42 | 0.42 | 0.58 | **0.51** | 0.0081 | 0.5052 | 0.5001 | 0.5006 | SSM |
| FitCons2 | 0.00 | 4.6549 | 0.49 | 0.51 | 0.59 | 0.41 | 0.41 | 0.59 | 0.50 | 0.0050 | 0.5000 | NA | 0.5000 | SSM |
| FunSeq2 | 0.00 | 1.1356 | 0.49 | 0.52 | 0.22 | 0.78 | 0.77 | 0.23 | 0.50 | 0.0120 | 0.5046 | 0.5005 | NA | SSM |
| LINSIGHT | 0.00 | 3.5183 | 0.50 | 0.51 | 0.53 | 0.47 | 0.46 | 0.54 | **0.51** | 0.0097 | 0.5058 | 0.5007 | 0.5006 | SSM |

*Note:* Best-threshold, the threshold corresponding to the best sum of sensitivity and specificity; PPV, positive predictive value; NPV, negative predictive value; FPR, false positive rate; FNR, false negative rate; MCC, mathew correlation coefficient; AUC, area under the curve; hspr-threshold, high-specificity regional threshold; hspr-AUC, high-specificity regional area under the curve; hser-threshold, high-sensitivity regional threshold; hser-AUC, high-sensitivity regional area under the curve; NA, not available; SM, supervised model; UM, unsupervised model; SSM, semi-supervised model. Top three methods of every measure are represented by bold text.

**Table S6.** Performance evaluation based on *de novo* mutation dataset.

| Method | Threshold | ASD | Sibling | OR | %95CI | %95CI | P value | Group |
| --- | --- | --- | --- | --- | --- | --- | --- | --- |
| CADD | 10.0000 | 9,498 | 9,232 | 1.0288 | 0.9997 | 1.0588 | 0.0528 | Pathogenic threshold = 10 |
| CDTS | 10.0000 | 10,861 | 10,533 | 1.0311 | 1.0038 | 1.0593 | 0.0254 | Pathogenic threshold = 10 |
| CScape | 10.0000 | 6,792 | 6,594 | 1.0300 | 0.9956 | 1.0657 | 0.0886 | Pathogenic threshold = 10 |
| DANN | 10.0000 | 9,478 | 9,005 | 1.0525 | 1.0225 | 1.0834 | 0.0005 | Pathogenic threshold = 10 |
| DVAR | 10.0000 | 11,255 | 10,972 | 1.0258 | 0.9991 | 1.0532 | 0.0586 | Pathogenic threshold = 10 |
| Eigen_PC | 10.0000 | 13,658 | 13,269 | 1.0293 | 1.0049 | 1.0543 | 0.0181 | Pathogenic threshold = 10 |
| Eigen | 10.0000 | 11,128 | 10,820 | 1.0285 | 1.0015 | 1.0561 | 0.0382 | Pathogenic threshold = 10 |
| FATHMM-MKL | 10.0000 | 9,333 | 9,111 | 1.0244 | 0.9951 | 1.0545 | 0.1037 | Pathogenic threshold = 10 |
| FATHMM-XF | 10.0000 | 10,358 | 10,233 | 1.0122 | 0.9848 | 1.0403 | 0.3875 | Pathogenic threshold = 10 |
| FIRE | 10.0000 | 11,593 | 11,449 | 1.0126 | 0.9867 | 1.0392 | 0.3462 | Pathogenic threshold = 10 |
| fitCons | 10.0000 | 13,966 | 13,541 | 1.0314 | 1.0072 | 1.0561 | 0.0106 | Pathogenic threshold = 10 |
| FitCons2 | 10.0000 | 11,925 | 11,533 | 1.0340 | 1.0078 | 1.0609 | 0.0107 | Pathogenic threshold = 10 |
| FunSeq2 | 10.0000 | 12,185 | 11,910 | 1.0231 | 0.9975 | 1.0493 | 0.0775 | Pathogenic threshold = 10 |
| GenoCanyon | 10.0000 | 13,418 | 12,980 | 1.0337 | 1.0090 | 1.0591 | 0.0072 | Pathogenic threshold = 10 |
| LINSIGHT | 10.0000 | 12,064 | 11,655 | 1.0351 | 1.0090 | 1.0619 | 0.0081 | Pathogenic threshold = 10 |
| ncER | 10.0000 | 9,183 | 8,901 | 1.0317 | 1.0019 | 1.0623 | 0.0367 | Pathogenic threshold = 10 |
| Orion | 10.0000 | 12,205 | 11,571 | 1.0548 | 1.0282 | 1.0821 | 0.0000 | Pathogenic threshold = 10 |
| PAFA | 10.0000 | 12,090 | 11,747 | 1.0292 | 1.0033 | 1.0558 | 0.0267 | Pathogenic threshold = 10 |
| ReMM | 10.0000 | 10,136 | 9,856 | 1.0284 | 1.0002 | 1.0574 | 0.0485 | Pathogenic threshold = 10 |
| regBase_REG | 10.0000 | 12,594 | 12,363 | 1.0187 | 0.9936 | 1.0444 | 0.1454 | Pathogenic threshold = 10 |
| regBase_CAN | 10.0000 | 9,430 | 9,425 | 1.0005 | 0.9723 | 1.0296 | 0.9768 | Pathogenic threshold = 10 |
| regBase_PAT | 10.0000 | 6,460 | 6,405 | 1.0086 | 0.9742 | 1.0442 | 0.6340 | Pathogenic threshold = 10 |
| DIVAN_TSS | 10.0000 | 12,357 | 12,240 | 1.0096 | 0.9846 | 1.0352 | 0.4595 | Pathogenic threshold = 10 |
| DIVAN_REGION | 10.0000 | 11,840 | 11,770 | 1.0059 | 0.9805 | 1.0320 | 0.6534 | Pathogenic threshold = 10 |
| CADD | 15.0000 | 2,769 | 2,589 | 1.0695 | 1.0134 | 1.1288 | 0.0145 | Pathogenic threshold = 15 |
| CDTS | 15.0000 | 3,192 | 3,005 | 1.0622 | 1.0103 | 1.1169 | 0.0181 | Pathogenic threshold = 15 |
| CScape | 15.0000 | 1,957 | 1,982 | 0.9874 | 0.9271 | 1.0516 | 0.7022 | Pathogenic threshold = 15 |
| DANN | 15.0000 | 2,595 | 2,527 | 1.0269 | 0.9718 | 1.0852 | 0.3492 | Pathogenic threshold = 15 |
| DVAR | 15.0000 | 2,702 | 2,600 | 1.0392 | 0.9844 | 1.0971 | 0.1654 | Pathogenic threshold = 15 |
| Eigen_PC | 15.0000 | 4,691 | 4,516 | 1.0388 | 0.9970 | 1.0823 | 0.0698 | Pathogenic threshold = 15 |
| Eigen | 15.0000 | 3,685 | 3,566 | 1.0334 | 0.9866 | 1.0824 | 0.1658 | Pathogenic threshold = 15 |
| FATHMM-MKL | 15.0000 | 2,590 | 2,557 | 1.0129 | 0.9587 | 1.0702 | 0.6556 | Pathogenic threshold = 15 |
| FATHMM-XF | 15.0000 | 3,259 | 3,129 | 1.0415 | 0.9914 | 1.0943 | 0.1065 | Pathogenic threshold = 15 |
| FIRE | 15.0000 | 3,858 | 3,826 | 1.0084 | 0.9640 | 1.0548 | 0.7236 | Pathogenic threshold = 15 |
| fitCons | 15.0000 | 2,971 | 2,776 | 1.0702 | 1.0159 | 1.1275 | 0.0105 | Pathogenic threshold = 15 |
| FitCons2 | 15.0000 | 2,657 | 2,468 | 1.0766 | 1.0188 | 1.1377 | 0.0086 | Pathogenic threshold = 15 |
| FunSeq2 | 15.0000 | 3,933 | 3,775 | 1.0419 | 0.9961 | 1.0897 | 0.0737 | Pathogenic threshold = 15 |
| GenoCanyon | 15.0000 | 4,349 | 4,241 | 1.0255 | 0.9828 | 1.0700 | 0.2483 | Pathogenic threshold = 15 |
| LINSIGHT | 15.0000 | 3,666 | 3,463 | 1.0586 | 1.0103 | 1.1093 | 0.0167 | Pathogenic threshold = 15 |
| ncER | 15.0000 | 2,847 | 2,806 | 1.0146 | 0.9627 | 1.0693 | 0.5947 | Pathogenic threshold = 15 |
| Orion | 15.0000 | 3,604 | 3,429 | 1.0510 | 1.0027 | 1.1017 | 0.0380 | Pathogenic threshold = 15 |
| PAFA | 15.0000 | 4,132 | 3,896 | 1.0606 | 1.0149 | 1.1083 | 0.0087 | Pathogenic threshold = 15 |
| ReMM | 15.0000 | 2,837 | 2,719 | 1.0434 | 0.9896 | 1.1002 | 0.1165 | Pathogenic threshold = 15 |
| regBase_REG | 15.0000 | 4,087 | 3,947 | 1.0355 | 0.9909 | 1.0820 | 0.1210 | Pathogenic threshold = 15 |
| regBase_CAN | 15.0000 | 1,971 | 2,088 | 0.9440 | 0.8872 | 1.0044 | 0.0686 | Pathogenic threshold = 15 |
| regBase_PAT | 15.0000 | 1,548 | 1,540 | 1.0052 | 0.9361 | 1.0794 | 0.8998 | Pathogenic threshold = 15 |
| DIVAN_TSS | 15.0000 | 3,736 | 3,894 | 0.9594 | 0.9171 | 1.0037 | 0.0723 | Pathogenic threshold = 15 |
| DIVAN_REGION | 15.0000 | 3,683 | 3,611 | 1.0199 | 0.9739 | 1.0681 | 0.4058 | Pathogenic threshold = 15 |
| CADD | 20.0000 | 453 | 401 | 1.1297 | 0.9854 | 1.2954 | 0.0809 | Pathogenic threshold = 20 |
| CDTS | 20.0000 | 1,077 | 952 | 1.1313 | 1.0359 | 1.2357 | 0.0059 | Pathogenic threshold = 20 |
| CScape | 20.0000 | 598 | 667 | 0.8966 | 0.8015 | 1.0027 | 0.0558 | Pathogenic threshold = 20 |
| DANN | 20.0000 | 383 | 384 | 0.9974 | 0.8635 | 1.1521 | 1.0000 | Pathogenic threshold = 20 |
| DVAR | 20.0000 | 727 | 710 | 1.0239 | 0.9221 | 1.1371 | 0.6730 | Pathogenic threshold = 20 |
| Eigen_PC | 20.0000 | 1,595 | 1,489 | 1.0712 | 0.9975 | 1.1504 | 0.0586 | Pathogenic threshold = 20 |
| Eigen | 20.0000 | 1,224 | 1,171 | 1.0453 | 0.9640 | 1.1334 | 0.2880 | Pathogenic threshold = 20 |
| FATHMM-MKL | 20.0000 | 653 | 662 | 0.9864 | 0.8840 | 1.1007 | 0.8254 | Pathogenic threshold = 20 |
| FATHMM-XF | 20.0000 | 1,028 | 1,015 | 1.0128 | 0.9278 | 1.1057 | 0.7906 | Pathogenic threshold = 20 |
| FIRE | 20.0000 | 1,242 | 1,245 | 0.9976 | 0.9214 | 1.0800 | 0.9680 | Pathogenic threshold = 20 |
| fitCons | 20.0000 | 62 | 69 | 0.8986 | 0.6272 | 1.2849 | 0.6003 | Pathogenic threshold = 20 |
| FitCons2 | 20.0000 | 71 | 66 | 1.0758 | 0.7585 | 1.5275 | 0.7327 | Pathogenic threshold = 20 |
| FunSeq2 | 20.0000 | 1,335 | 1,335 | 1.0000 | 0.9262 | 1.0796 | 1.0000 | Pathogenic threshold = 20 |
| GenoCanyon | 20.0000 | 1,422 | 1,342 | 1.0596 | 0.9828 | 1.1425 | 0.1329 | Pathogenic threshold = 20 |
| LINSIGHT | 20.0000 | 1,102 | 1,082 | 1.0185 | 0.9357 | 1.1086 | 0.6843 | Pathogenic threshold = 20 |
| ncER | 20.0000 | 910 | 942 | 0.9660 | 0.8809 | 1.0593 | 0.4713 | Pathogenic threshold = 20 |
| Orion | 20.0000 | 1,138 | 1,019 | 1.1168 | 1.0254 | 1.2165 | 0.0110 | Pathogenic threshold = 20 |
| PAFA | 20.0000 | 1,576 | 1,444 | 1.0914 | 1.0156 | 1.1730 | 0.0171 | Pathogenic threshold = 20 |
| ReMM | 20.0000 | 727 | 689 | 1.0552 | 0.9494 | 1.1727 | 0.3255 | Pathogenic threshold = 20 |
| regBase_REG | 20.0000 | 1,331 | 1,265 | 1.0522 | 0.9735 | 1.1372 | 0.2020 | Pathogenic threshold = 20 |
| regBase_CAN | 20.0000 | 584 | 640 | 0.9125 | 0.8143 | 1.0224 | 0.1159 | Pathogenic threshold = 20 |
| regBase_PAT | 20.0000 | 482 | 479 | 1.0063 | 0.8849 | 1.1443 | 0.9486 | Pathogenic threshold = 20 |
| DIVAN_TSS | 20.0000 | 1,136 | 1,206 | 0.9420 | 0.8679 | 1.0223 | 0.1539 | Pathogenic threshold = 20 |
| DIVAN_REGION | 20.0000 | 1,132 | 1,086 | 1.0424 | 0.9582 | 1.1339 | 0.3393 | Pathogenic threshold = 20 |
| CADD | 25.0000 | 2 | 1 | 2.0000 | 0.1041 | 117.9944 | 1.0000 | Pathogenic threshold = 25 |
| CDTS | 25.0000 | 362 | 332 | 1.0904 | 0.9369 | 1.2693 | 0.2710 | Pathogenic threshold = 25 |
| CScape | 25.0000 | 211 | 231 | 0.9134 | 0.7543 | 1.1057 | 0.3662 | Pathogenic threshold = 25 |
| DANN | 25.0000 | 4 | 3 | 1.3333 | 0.2256 | 9.1022 | 1.0000 | Pathogenic threshold = 25 |
| DVAR | 25.0000 | 252 | 235 | 1.0723 | 0.8941 | 1.2864 | 0.4685 | Pathogenic threshold = 25 |
| Eigen_PC | 25.0000 | 553 | 516 | 1.0717 | 0.9488 | 1.2107 | 0.2709 | Pathogenic threshold = 25 |
| Eigen | 25.0000 | 421 | 404 | 1.0421 | 0.9069 | 1.1975 | 0.5775 | Pathogenic threshold = 25 |
| FATHMM-MKL | 25.0000 | 121 | 157 | 0.7707 | 0.6029 | 0.9832 | 0.0356 | Pathogenic threshold = 25 |
| FATHMM-XF | 25.0000 | 279 | 301 | 0.9269 | 0.7847 | 1.0945 | 0.3832 | Pathogenic threshold = 25 |
| FIRE | 25.0000 | 385 | 397 | 0.9698 | 0.8407 | 1.1186 | 0.6941 | Pathogenic threshold = 25 |
| fitCons | 25.0000 | 14 | 14 | 1.0000 | 0.4419 | 2.2630 | 1.0000 | Pathogenic threshold = 25 |
| FitCons2 | 25.0000 | 25 | 19 | 1.3158 | 0.6959 | 2.5274 | 0.4514 | Pathogenic threshold = 25 |
| FunSeq2 | 25.0000 | 466 | 454 | 1.0264 | 0.9000 | 1.1706 | 0.7169 | Pathogenic threshold = 25 |
| GenoCanyon | 25.0000 | 474 | 464 | 1.0216 | 0.8969 | 1.1636 | 0.7689 | Pathogenic threshold = 25 |
| LINSIGHT | 25.0000 | 393 | 360 | 1.0917 | 0.9438 | 1.2630 | 0.2435 | Pathogenic threshold = 25 |
| ncER | 25.0000 | 264 | 276 | 0.9565 | 0.8049 | 1.1365 | 0.6360 | Pathogenic threshold = 25 |
| Orion | 25.0000 | 332 | 303 | 1.0957 | 0.9349 | 1.2846 | 0.2665 | Pathogenic threshold = 25 |
| PAFA | 25.0000 | 844 | 782 | 1.0793 | 0.9781 | 1.1911 | 0.1303 | Pathogenic threshold = 25 |
| ReMM | 25.0000 | 244 | 214 | 1.1402 | 0.9451 | 1.3764 | 0.1753 | Pathogenic threshold = 25 |
| regBase_REG | 25.0000 | 419 | 385 | 1.0883 | 0.9454 | 1.2531 | 0.2445 | Pathogenic threshold = 25 |
| regBase_CAN | 25.0000 | 191 | 191 | 1.0000 | 0.8140 | 1.2286 | 1.0000 | Pathogenic threshold = 25 |
| regBase_PAT | 25.0000 | 138 | 133 | 1.0376 | 0.8117 | 1.3267 | 0.8081 | Pathogenic threshold = 25 |
| DIVAN_TSS | 25.0000 | 355 | 362 | 0.9807 | 0.8447 | 1.1384 | 0.8227 | Pathogenic threshold = 25 |
| DIVAN_REGION | 25.0000 | 358 | 334 | 1.0719 | 0.9208 | 1.2480 | 0.3820 | Pathogenic threshold = 25 |
| CADD | 30.0000 | 1 | 0 | 1.0000 | 0.0256 | 1.0000 | 1.0000 | Pathogenic threshold = 30 |
| CDTS | 30.0000 | 119 | 90 | 1.3222 | 0.9971 | 1.7585 | 0.0525 | Pathogenic threshold = 30 |
| CScape | 30.0000 | 76 | 83 | 0.9157 | 0.6619 | 1.2653 | 0.6343 | Pathogenic threshold = 30 |
| DANN | 30.0000 | 2 | 0 | 1.0000 | 0.1878 | 1.0000 | 0.5000 | Pathogenic threshold = 30 |
| DVAR | 30.0000 | 65 | 74 | 0.8784 | 0.6196 | 1.2426 | 0.4976 | Pathogenic threshold = 30 |
| Eigen_PC | 30.0000 | 206 | 174 | 1.1839 | 0.9628 | 1.4571 | 0.1117 | Pathogenic threshold = 30 |
| Eigen | 30.0000 | 155 | 145 | 1.0690 | 0.8468 | 1.3500 | 0.6034 | Pathogenic threshold = 30 |
| FATHMM-MKL | 30.0000 | 45 | 62 | 0.7258 | 0.4832 | 1.0827 | 0.1215 | Pathogenic threshold = 30 |
| FATHMM-XF | 30.0000 | 74 | 67 | 1.1045 | 0.7828 | 1.5607 | 0.6135 | Pathogenic threshold = 30 |
| FIRE | 30.0000 | 144 | 138 | 1.0435 | 0.8204 | 1.3276 | 0.7660 | Pathogenic threshold = 30 |
| fitCons | 30.0000 | 3 | 4 | 0.7500 | 0.1099 | 4.4333 | 1.0000 | Pathogenic threshold = 30 |
| FitCons2 | 30.0000 | 24 | 18 | 1.3333 | 0.6938 | 2.6074 | 0.4408 | Pathogenic threshold = 30 |
| FunSeq2 | 30.0000 | 170 | 137 | 1.2409 | 0.9850 | 1.5656 | 0.0676 | Pathogenic threshold = 30 |
| GenoCanyon | 30.0000 | 306 | 269 | 1.1375 | 0.9625 | 1.3451 | 0.1332 | Pathogenic threshold = 30 |
| LINSIGHT | 30.0000 | 136 | 119 | 1.1429 | 0.8870 | 1.4742 | 0.3164 | Pathogenic threshold = 30 |
| ncER | 30.0000 | 47 | 61 | 0.7705 | 0.5152 | 1.1459 | 0.2108 | Pathogenic threshold = 30 |
| Orion | 30.0000 | 97 | 96 | 1.0104 | 0.7541 | 1.3541 | 1.0000 | Pathogenic threshold = 30 |
| PAFA | 30.0000 | 844 | 782 | 1.0793 | 0.9781 | 1.1911 | 0.1303 | Pathogenic threshold = 30 |
| ReMM | 30.0000 | 84 | 69 | 1.2174 | 0.8748 | 1.6990 | 0.2576 | Pathogenic threshold = 30 |
| regBase_REG | 30.0000 | 137 | 121 | 1.1322 | 0.8801 | 1.4580 | 0.3504 | Pathogenic threshold = 30 |
| regBase_CAN | 30.0000 | 53 | 48 | 1.1042 | 0.7330 | 1.6671 | 0.6908 | Pathogenic threshold = 30 |
| regBase_PAT | 30.0000 | 63 | 64 | 0.9844 | 0.6840 | 1.4163 | 1.0000 | Pathogenic threshold = 30 |
| DIVAN_TSS | 30.0000 | 113 | 105 | 1.0762 | 0.8178 | 1.4173 | 0.6355 | Pathogenic threshold = 30 |
| DIVAN_REGION | 30.0000 | 117 | 108 | 1.0833 | 0.8268 | 1.4205 | 0.5939 | Pathogenic threshold = 30 |
| CADD | 35.0000 | 0 | 0 | 1.0000 | 0.0000 | 1.0000 | 1.0000 | Pathogenic threshold = 35 |
| CDTS | 35.0000 | 32 | 20 | 1.6000 | 0.8875 | 2.9518 | 0.1263 | Pathogenic threshold = 35 |
| CScape | 35.0000 | 11 | 13 | 0.8462 | 0.3432 | 2.0468 | 0.8388 | Pathogenic threshold = 35 |
| DANN | 35.0000 | 1 | 0 | 1.0000 | 0.0256 | 1.0000 | 1.0000 | Pathogenic threshold = 35 |
| DVAR | 35.0000 | 32 | 30 | 1.0667 | 0.6277 | 1.8171 | 0.8991 | Pathogenic threshold = 35 |
| Eigen_PC | 35.0000 | 71 | 65 | 1.0923 | 0.7692 | 1.5535 | 0.6683 | Pathogenic threshold = 35 |
| Eigen | 35.0000 | 57 | 52 | 1.0962 | 0.7394 | 1.6281 | 0.7018 | Pathogenic threshold = 35 |
| FATHMM-MKL | 35.0000 | 33 | 46 | 0.7174 | 0.4444 | 1.1465 | 0.1766 | Pathogenic threshold = 35 |
| FATHMM-XF | 35.0000 | 14 | 12 | 1.1667 | 0.5008 | 2.7612 | 0.8450 | Pathogenic threshold = 35 |
| FIRE | 35.0000 | 57 | 39 | 1.4615 | 0.9557 | 2.2560 | 0.0822 | Pathogenic threshold = 35 |
| fitCons | 35.0000 | 1 | 3 | 0.3333 | 0.0063 | 4.1514 | 0.6250 | Pathogenic threshold = 35 |
| FitCons2 | 35.0000 | 23 | 18 | 1.2778 | 0.6597 | 2.5126 | 0.5327 | Pathogenic threshold = 35 |
| FunSeq2 | 35.0000 | 45 | 44 | 1.0227 | 0.6598 | 1.5862 | 1.0000 | Pathogenic threshold = 35 |
| GenoCanyon | 35.0000 | 306 | 269 | 1.1375 | 0.9625 | 1.3451 | 0.1332 | Pathogenic threshold = 35 |
| LINSIGHT | 35.0000 | 42 | 41 | 1.0244 | 0.6501 | 1.6152 | 1.0000 | Pathogenic threshold = 35 |
| ncER | 35.0000 | 2 | 10 | 0.2000 | 0.0213 | 0.9385 | 0.0386 | Pathogenic threshold = 35 |
| Orion | 35.0000 | 20 | 27 | 0.7407 | 0.3939 | 1.3709 | 0.3817 | Pathogenic threshold = 35 |
| PAFA | 35.0000 | 844 | 782 | 1.0793 | 0.9781 | 1.1911 | 0.1303 | Pathogenic threshold = 35 |
| ReMM | 35.0000 | 25 | 25 | 1.0000 | 0.5510 | 1.8147 | 1.0000 | Pathogenic threshold = 35 |
| regBase_REG | 35.0000 | 38 | 26 | 1.4615 | 0.8645 | 2.5070 | 0.1686 | Pathogenic threshold = 35 |
| regBase_CAN | 35.0000 | 17 | 24 | 0.7083 | 0.3572 | 1.3748 | 0.3489 | Pathogenic threshold = 35 |
| regBase_PAT | 35.0000 | 34 | 34 | 1.0000 | 0.6031 | 1.6581 | 1.0000 | Pathogenic threshold = 35 |
| DIVAN_TSS | 35.0000 | 36 | 36 | 1.0000 | 0.6123 | 1.6333 | 1.0000 | Pathogenic threshold = 35 |
| DIVAN_REGION | 35.0000 | 41 | 38 | 1.0789 | 0.6769 | 1.7238 | 0.8221 | Pathogenic threshold = 35 |
| CADD | 22.2285 | 50 | 35 | 1.4286 | 0.9092 | 2.2672 | 0.1284 | Top 50 DNMs in ASD |
| CDTS | 32.7426 | 50 | 40 | 1.2500 | 0.8083 | 1.9443 | 0.3428 | Top 50 DNMs in ASD |
| CScape | 31.0251 | 50 | 59 | 0.8475 | 0.5696 | 1.2566 | 0.4437 | Top 50 DNMs in ASD |
| DANN | 21.8926 | 50 | 47 | 1.0638 | 0.7000 | 1.6192 | 0.8392 | Top 50 DNMs in ASD |
| DIVAN_REGION | 19.0293 | 51 | 42 | 1.2143 | 0.7913 | 1.8724 | 0.4069 | Top 50 DNMs in ASD |
| DIVAN_TSS | 18.8821 | 51 | 49 | 1.0408 | 0.6893 | 1.5730 | 0.9204 | Top 50 DNMs in ASD |
| DVAR | 31.3630 | 60 | 68 | 0.8824 | 0.6130 | 1.2673 | 0.5363 | Top 50 DNMs in ASD |
| Eigen | 35.9826 | 50 | 38 | 1.3158 | 0.8458 | 2.0620 | 0.2408 | Top 50 DNMs in ASD |
| Eigen_PC | 36.7700 | 50 | 48 | 1.0417 | 0.6868 | 1.5813 | 0.9196 | Top 50 DNMs in ASD |
| FATHMM-MKL | 28.7705 | 50 | 72 | 0.6944 | 0.4742 | 1.0101 | 0.0568 | Top 50 DNMs in ASD |
| FATHMM-XF | 31.8651 | 50 | 43 | 1.1628 | 0.7580 | 1.7905 | 0.5341 | Top 50 DNMs in ASD |
| FIRE | 35.9983 | 57 | 39 | 1.4615 | 0.9557 | 2.2560 | 0.0822 | Top 50 DNMs in ASD |
| fitCons | 20.4645 | 50 | 50 | 1.0000 | 0.6620 | 1.5105 | 1.0000 | Top 50 DNMs in ASD |
| FitCons2 | 20.2120 | 71 | 66 | 1.0758 | 0.7585 | 1.5275 | 0.7327 | Top 50 DNMs in ASD |
| FunSeq2 | 34.8172 | 50 | 45 | 1.1111 | 0.7278 | 1.7006 | 0.6817 | Top 50 DNMs in ASD |
| GenoCanyon | 99.3328 | 306 | 269 | 1.1375 | 0.9625 | 1.3451 | 0.1332 | Top 50 DNMs in ASD |
| LINSIGHT | 34.3931 | 50 | 45 | 1.1111 | 0.7278 | 1.7006 | 0.6817 | Top 50 DNMs in ASD |
| ncER | 29.7367 | 50 | 68 | 0.7353 | 0.4999 | 1.0751 | 0.1172 | Top 50 DNMs in ASD |
| Orion | 31.9212 | 50 | 59 | 0.8475 | 0.5696 | 1.2566 | 0.4437 | Top 50 DNMs in ASD |
| PAFA | 99.3328 | 844 | 782 | 1.0793 | 0.9781 | 1.1911 | 0.1303 | Top 50 DNMs in ASD |
| regBase_REG | 34.1327 | 50 | 39 | 1.2821 | 0.8266 | 2.0014 | 0.2891 | Top 50 DNMs in ASD |
| regBase_CAN | 30.1987 | 50 | 46 | 1.0870 | 0.7136 | 1.6589 | 0.7596 | Top 50 DNMs in ASD |
| regBase_PAT | 31.2330 | 50 | 53 | 0.9434 | 0.6280 | 1.4153 | 0.8439 | Top 50 DNMs in ASD |
| ReMM | 34.4589 | 53 | 39 | 1.3590 | 0.8819 | 2.1106 | 0.1750 | Top 50 DNMs in ASD |
| CADD | 21.8455 | 100 | 76 | 1.3158 | 0.9667 | 1.7971 | 0.0827 | Top 100 DNMs in ASD |
| CDTS | 30.4395 | 100 | 79 | 1.2658 | 0.9330 | 1.7224 | 0.1347 | Top 100 DNMs in ASD |
| CScape | 29.5308 | 100 | 97 | 1.0309 | 0.7718 | 1.3775 | 0.8867 | Top 100 DNMs in ASD |
| DANN | 21.5279 | 100 | 105 | 0.9524 | 0.7169 | 1.2645 | 0.7800 | Top 100 DNMs in ASD |
| DIVAN_REGION | 30.9855 | 101 | 89 | 1.1348 | 0.8450 | 1.5264 | 0.4249 | Top 100 DNMs in ASD |
| DIVAN_TSS | 30.9182 | 100 | 81 | 1.2346 | 0.9118 | 1.6759 | 0.1808 | Top 100 DNMs in ASD |
| DVAR | 28.7393 | 106 | 110 | 0.9636 | 0.7310 | 1.2699 | 0.8383 | Top 100 DNMs in ASD |
| Eigen | 32.3566 | 100 | 93 | 1.0753 | 0.8026 | 1.4418 | 0.6659 | Top 100 DNMs in ASD |
| Eigen_PC | 33.4089 | 100 | 95 | 1.0526 | 0.7869 | 1.4089 | 0.7746 | Top 100 DNMs in ASD |
| FATHMM-MKL | 25.5653 | 100 | 131 | 0.7634 | 0.5825 | 0.9979 | 0.0482 | Top 100 DNMs in ASD |
| FATHMM-XF | 28.9296 | 100 | 96 | 1.0417 | 0.7793 | 1.3930 | 0.8304 | Top 100 DNMs in ASD |
| FIRE | 32.5824 | 100 | 77 | 1.2987 | 0.9552 | 1.7715 | 0.0979 | Top 100 DNMs in ASD |
| fitCons | 19.2303 | 120 | 115 | 1.0435 | 0.8012 | 1.3595 | 0.7942 | Top 100 DNMs in ASD |
| FitCons2 | 19.5952 | 119 | 119 | 1.0000 | 0.7690 | 1.3003 | 1.0000 | Top 100 DNMs in ASD |
| FunSeq2 | 32.3758 | 100 | 75 | 1.3333 | 0.9785 | 1.8235 | 0.0693 | Top 100 DNMs in ASD |
| GenoCanyon | 99.3328 | 306 | 269 | 1.1375 | 0.9625 | 1.3451 | 0.1332 | Top 100 DNMs in ASD |
| LINSIGHT | 31.2790 | 100 | 90 | 1.1111 | 0.8274 | 1.4940 | 0.5139 | Top 100 DNMs in ASD |
| ncER | 28.1597 | 100 | 112 | 0.8929 | 0.6749 | 1.1798 | 0.4500 | Top 100 DNMs in ASD |
| Orion | 29.8353 | 100 | 101 | 0.9901 | 0.7433 | 1.3186 | 1.0000 | Top 100 DNMs in ASD |
| PAFA | 99.3328 | 844 | 782 | 1.0793 | 0.9781 | 1.1911 | 0.1303 | Top 100 DNMs in ASD |
| regBase_REG | 31.4782 | 100 | 85 | 1.1765 | 0.8722 | 1.5899 | 0.3033 | Top 100 DNMs in ASD |
| regBase_CAN | 27.6023 | 100 | 81 | 1.2346 | 0.9118 | 1.6759 | 0.1808 | Top 100 DNMs in ASD |
| regBase_PAT | 26.7619 | 100 | 98 | 1.0204 | 0.7645 | 1.3623 | 0.9434 | Top 100 DNMs in ASD |
| ReMM | 29.0541 | 132 | 106 | 1.2453 | 0.9571 | 1.6235 | 0.1049 | Top 100 DNMs in ASD |
| CADD | 21.6241 | 150 | 100 | 1.5000 | 1.1569 | 1.9517 | 0.0019 | Top 150 DNMs in ASD |
| CDTS | 28.9781 | 150 | 123 | 1.2195 | 0.9544 | 1.5607 | 0.1154 | Top 150 DNMs in ASD |
| CScape | 27.6840 | 150 | 151 | 0.9934 | 0.7872 | 1.2536 | 1.0000 | Top 150 DNMs in ASD |
| DANN | 21.1465 | 150 | 167 | 0.8982 | 0.7156 | 1.1266 | 0.3689 | Top 150 DNMs in ASD |
| DIVAN_REGION | 28.4792 | 151 | 153 | 0.9869 | 0.7830 | 1.2439 | 0.9543 | Top 150 DNMs in ASD |
| DIVAN_TSS | 28.8467 | 152 | 146 | 1.0411 | 0.8241 | 1.3156 | 0.7721 | Top 150 DNMs in ASD |
| DVAR | 26.8063 | 167 | 162 | 1.0309 | 0.8255 | 1.2876 | 0.8255 | Top 150 DNMs in ASD |
| Eigen | 30.2956 | 150 | 131 | 1.1450 | 0.8997 | 1.4587 | 0.2829 | Top 150 DNMs in ASD |
| Eigen_PC | 31.5039 | 150 | 127 | 1.1811 | 0.9263 | 1.5080 | 0.1861 | Top 150 DNMs in ASD |
| FATHMM-MKL | 24.6066 | 151 | 179 | 0.8436 | 0.6748 | 1.0535 | 0.1371 | Top 150 DNMs in ASD |
| FATHMM-XF | 27.4740 | 150 | 140 | 1.0714 | 0.8453 | 1.3587 | 0.5972 | Top 150 DNMs in ASD |
| FIRE | 29.6693 | 160 | 161 | 0.9938 | 0.7935 | 1.2446 | 1.0000 | Top 150 DNMs in ASD |
| fitCons | 19.1281 | 306 | 280 | 1.0929 | 0.9263 | 1.2898 | 0.3017 | Top 150 DNMs in ASD |
| FitCons2 | 19.4405 | 372 | 342 | 1.0877 | 0.9366 | 1.2635 | 0.2778 | Top 150 DNMs in ASD |
| FunSeq2 | 30.5943 | 150 | 121 | 1.2397 | 0.9691 | 1.5884 | 0.0888 | Top 150 DNMs in ASD |
| GenoCanyon | 99.3328 | 306 | 269 | 1.1375 | 0.9625 | 1.3451 | 0.1332 | Top 150 DNMs in ASD |
| LINSIGHT | 29.5931 | 150 | 132 | 1.1364 | 0.8933 | 1.4469 | 0.3114 | Top 150 DNMs in ASD |
| ncER | 26.6932 | 150 | 176 | 0.8523 | 0.6809 | 1.0657 | 0.1661 | Top 150 DNMs in ASD |
| Orion | 28.3423 | 150 | 143 | 1.0490 | 0.8286 | 1.3284 | 0.7260 | Top 150 DNMs in ASD |
| PAFA | 99.3328 | 844 | 782 | 1.0793 | 0.9781 | 1.1911 | 0.1303 | Top 150 DNMs in ASD |
| regBase_REG | 29.5488 | 150 | 137 | 1.0949 | 0.8627 | 1.3905 | 0.4788 | Top 150 DNMs in ASD |
| regBase_CAN | 25.6810 | 150 | 152 | 0.9868 | 0.7823 | 1.2448 | 0.9541 | Top 150 DNMs in ASD |
| regBase_PAT | 24.6028 | 150 | 144 | 1.0417 | 0.8232 | 1.3185 | 0.7706 | Top 150 DNMs in ASD |
| ReMM | 27.4452 | 169 | 140 | 1.2071 | 0.9592 | 1.5212 | 0.1110 | Top 150 DNMs in ASD |
| CADD | 21.2419 | 200 | 149 | 1.3423 | 1.0803 | 1.6708 | 0.0074 | Top 200 DNMs in ASD |
| CDTS | 27.9946 | 200 | 165 | 1.2121 | 0.9814 | 1.4988 | 0.0750 | Top 200 DNMs in ASD |
| CScape | 25.2729 | 200 | 223 | 0.8969 | 0.7373 | 1.0904 | 0.2848 | Top 200 DNMs in ASD |
| DANN | 20.8830 | 200 | 214 | 0.9346 | 0.7668 | 1.1386 | 0.5229 | Top 200 DNMs in ASD |
| DIVAN_REGION | 27.3773 | 203 | 205 | 0.9902 | 0.8115 | 1.2083 | 0.9605 | Top 200 DNMs in ASD |
| DIVAN_TSS | 27.4981 | 200 | 195 | 1.0256 | 0.8378 | 1.2558 | 0.8405 | Top 200 DNMs in ASD |
| DVAR | 25.9737 | 205 | 191 | 1.0733 | 0.8770 | 1.3141 | 0.5136 | Top 200 DNMs in ASD |
| Eigen | 28.5822 | 200 | 194 | 1.0309 | 0.8419 | 1.2626 | 0.8012 | Top 200 DNMs in ASD |
| Eigen_PC | 30.1946 | 200 | 165 | 1.2121 | 0.9814 | 1.4988 | 0.0750 | Top 200 DNMs in ASD |
| FATHMM-MKL | 23.7084 | 200 | 245 | 0.8163 | 0.6738 | 0.9880 | 0.0369 | Top 200 DNMs in ASD |
| FATHMM-XF | 26.2866 | 200 | 202 | 0.9901 | 0.8102 | 1.2099 | 0.9602 | Top 200 DNMs in ASD |
| FIRE | 28.5879 | 204 | 205 | 0.9951 | 0.8157 | 1.2139 | 1.0000 | Top 200 DNMs in ASD |
| fitCons | 19.1281 | 306 | 280 | 1.0929 | 0.9263 | 1.2898 | 0.3017 | Top 200 DNMs in ASD |
| FitCons2 | 19.4405 | 372 | 342 | 1.0877 | 0.9366 | 1.2635 | 0.2778 | Top 200 DNMs in ASD |
| FunSeq2 | 29.2104 | 201 | 177 | 1.1356 | 0.9232 | 1.3978 | 0.2368 | Top 200 DNMs in ASD |
| GenoCanyon | 99.3328 | 306 | 269 | 1.1375 | 0.9625 | 1.3451 | 0.1332 | Top 200 DNMs in ASD |
| LINSIGHT | 28.1606 | 200 | 187 | 1.0695 | 0.8718 | 1.3126 | 0.5419 | Top 200 DNMs in ASD |
| ncER | 25.9870 | 200 | 208 | 0.9615 | 0.7879 | 1.1732 | 0.7290 | Top 200 DNMs in ASD |
| Orion | 27.2464 | 200 | 184 | 1.0870 | 0.8853 | 1.3352 | 0.4440 | Top 200 DNMs in ASD |
| PAFA | 99.3328 | 844 | 782 | 1.0793 | 0.9781 | 1.1911 | 0.1303 | Top 200 DNMs in ASD |
| regBase_REG | 28.1902 | 200 | 181 | 1.1050 | 0.8992 | 1.3586 | 0.3565 | Top 200 DNMs in ASD |
| regBase_CAN | 24.7373 | 200 | 202 | 0.9901 | 0.8102 | 1.2099 | 0.9602 | Top 200 DNMs in ASD |
| regBase_PAT | 23.1036 | 200 | 198 | 1.0101 | 0.8257 | 1.2357 | 0.9600 | Top 200 DNMs in ASD |
| ReMM | 26.2316 | 211 | 178 | 1.1854 | 0.9664 | 1.4553 | 0.1046 | Top 200 DNMs in ASD |
| CADD | 21.0103 | 250 | 192 | 1.3021 | 1.0745 | 1.5798 | 0.0066 | Top 250 DNMs in ASD |
| CDTS | 26.8855 | 250 | 207 | 1.2077 | 1.0005 | 1.4591 | 0.0493 | Top 250 DNMs in ASD |
| CScape | 24.1744 | 250 | 273 | 0.9158 | 0.7683 | 1.0912 | 0.3361 | Top 250 DNMs in ASD |
| DANN | 20.6242 | 250 | 270 | 0.9259 | 0.7764 | 1.1038 | 0.4048 | Top 250 DNMs in ASD |
| DIVAN_REGION | 26.5827 | 252 | 234 | 1.0769 | 0.8978 | 1.2922 | 0.4407 | Top 250 DNMs in ASD |
| DIVAN_TSS | 26.3557 | 252 | 253 | 0.9960 | 0.8333 | 1.1906 | 1.0000 | Top 250 DNMs in ASD |
| DVAR | 25.0378 | 252 | 235 | 1.0723 | 0.8941 | 1.2864 | 0.4685 | Top 250 DNMs in ASD |
| Eigen | 27.6044 | 250 | 240 | 1.0417 | 0.8691 | 1.2488 | 0.6844 | Top 250 DNMs in ASD |
| Eigen_PC | 29.0694 | 250 | 220 | 1.1364 | 0.9442 | 1.3684 | 0.1809 | Top 250 DNMs in ASD |
| FATHMM-MKL | 23.1645 | 250 | 284 | 0.8803 | 0.7397 | 1.0471 | 0.1532 | Top 250 DNMs in ASD |
| FATHMM-XF | 25.4267 | 250 | 265 | 0.9434 | 0.7905 | 1.1256 | 0.5373 | Top 250 DNMs in ASD |
| FIRE | 27.2538 | 261 | 255 | 1.0235 | 0.8580 | 1.2211 | 0.8258 | Top 250 DNMs in ASD |
| fitCons | 19.1281 | 306 | 280 | 1.0929 | 0.9263 | 1.2898 | 0.3017 | Top 250 DNMs in ASD |
| FitCons2 | 19.4405 | 372 | 342 | 1.0877 | 0.9366 | 1.2635 | 0.2778 | Top 250 DNMs in ASD |
| FunSeq2 | 28.1725 | 250 | 237 | 1.0549 | 0.8796 | 1.2654 | 0.5866 | Top 250 DNMs in ASD |
| GenoCanyon | 99.3328 | 306 | 269 | 1.1375 | 0.9625 | 1.3451 | 0.1332 | Top 250 DNMs in ASD |
| LINSIGHT | 26.9984 | 250 | 242 | 1.0331 | 0.8622 | 1.2379 | 0.7524 | Top 250 DNMs in ASD |
| ncER | 25.2497 | 250 | 258 | 0.9690 | 0.8110 | 1.1576 | 0.7562 | Top 250 DNMs in ASD |
| Orion | 26.4218 | 250 | 216 | 1.1574 | 0.9609 | 1.3951 | 0.1262 | Top 250 DNMs in ASD |
| PAFA | 99.3328 | 844 | 782 | 1.0793 | 0.9781 | 1.1911 | 0.1303 | Top 250 DNMs in ASD |
| regBase_REG | 27.2604 | 250 | 220 | 1.1364 | 0.9442 | 1.3684 | 0.1809 | Top 250 DNMs in ASD |
| regBase_CAN | 23.7142 | 250 | 268 | 0.9328 | 0.7820 | 1.1125 | 0.4551 | Top 250 DNMs in ASD |
| regBase_PAT | 22.1223 | 250 | 258 | 0.9690 | 0.8110 | 1.1576 | 0.7562 | Top 250 DNMs in ASD |
| ReMM | 24.5162 | 281 | 249 | 1.1285 | 0.9481 | 1.3438 | 0.1781 | Top 250 DNMs in ASD |
| CADD | 20.6685 | 300 | 258 | 1.1628 | 0.9812 | 1.3787 | 0.0825 | Top 300 DNMs in ASD |
| CDTS | 26.1352 | 300 | 254 | 1.1811 | 0.9960 | 1.4015 | 0.0558 | Top 300 DNMs in ASD |
| CScape | 23.3983 | 300 | 322 | 0.9317 | 0.7934 | 1.0938 | 0.3998 | Top 300 DNMs in ASD |
| DANN | 20.3789 | 300 | 312 | 0.9615 | 0.8179 | 1.1303 | 0.6566 | Top 300 DNMs in ASD |
| DIVAN_REGION | 25.7736 | 305 | 281 | 1.0854 | 0.9200 | 1.2810 | 0.3421 | Top 300 DNMs in ASD |
| DIVAN_TSS | 25.6030 | 300 | 312 | 0.9615 | 0.8179 | 1.1303 | 0.6566 | Top 300 DNMs in ASD |
| DVAR | 24.1092 | 308 | 300 | 1.0267 | 0.8729 | 1.2076 | 0.7765 | Top 300 DNMs in ASD |
| Eigen | 26.5387 | 300 | 310 | 0.9677 | 0.8229 | 1.1379 | 0.7156 | Top 300 DNMs in ASD |
| Eigen_PC | 27.9394 | 300 | 275 | 1.0909 | 0.9231 | 1.2896 | 0.3169 | Top 300 DNMs in ASD |
| FATHMM-MKL | 22.6063 | 300 | 332 | 0.9036 | 0.7704 | 1.0595 | 0.2175 | Top 300 DNMs in ASD |
| FATHMM-XF | 24.6196 | 300 | 333 | 0.9009 | 0.7682 | 1.0562 | 0.2034 | Top 300 DNMs in ASD |
| FIRE | 26.5044 | 310 | 297 | 1.0438 | 0.8873 | 1.2280 | 0.6262 | Top 300 DNMs in ASD |
| fitCons | 19.1281 | 306 | 280 | 1.0929 | 0.9263 | 1.2898 | 0.3017 | Top 300 DNMs in ASD |
| FitCons2 | 19.4405 | 372 | 342 | 1.0877 | 0.9366 | 1.2635 | 0.2778 | Top 300 DNMs in ASD |
| FunSeq2 | 27.2648 | 300 | 281 | 1.0676 | 0.9043 | 1.2608 | 0.4552 | Top 300 DNMs in ASD |
| GenoCanyon | 99.3328 | 306 | 269 | 1.1375 | 0.9625 | 1.3451 | 0.1332 | Top 300 DNMs in ASD |
| LINSIGHT | 26.2336 | 300 | 279 | 1.0753 | 0.9105 | 1.2702 | 0.4059 | Top 300 DNMs in ASD |
| ncER | 24.6434 | 300 | 301 | 0.9967 | 0.8466 | 1.1734 | 1.0000 | Top 300 DNMs in ASD |
| Orion | 25.5904 | 300 | 267 | 1.1236 | 0.9496 | 1.3300 | 0.1789 | Top 300 DNMs in ASD |
| PAFA | 99.3328 | 844 | 782 | 1.0793 | 0.9781 | 1.1911 | 0.1303 | Top 300 DNMs in ASD |
| regBase_REG | 26.4005 | 300 | 278 | 1.0791 | 0.9136 | 1.2750 | 0.3824 | Top 300 DNMs in ASD |
| regBase_CAN | 22.9110 | 300 | 332 | 0.9036 | 0.7704 | 1.0595 | 0.2175 | Top 300 DNMs in ASD |
| regBase_PAT | 21.6681 | 300 | 302 | 0.9934 | 0.8439 | 1.1694 | 0.9675 | Top 300 DNMs in ASD |
| ReMM | 23.8832 | 322 | 277 | 1.1625 | 0.9869 | 1.3700 | 0.0721 | Top 300 DNMs in ASD |

*Note:* The OR, 95% confidence interval and *P* values were calculated by Poisson's ratio test. DNMs, *de novo* mutations.

**Table S7.** Performance evaluation based on experimentally validated de novo mutations from ASD

| Methods | Missing (%) | Best-threshold | PPV (%) | NPV (%) | FNR (%) | Sensitivity (%) | FPR (%) | Specificity (%) | Accuracy (%) | MCC | AUC | hspr-AUC | hser-AUC | Predicted model |
| --- | --- | --- | --- | --- | --- | --- | --- | --- | --- | --- | --- | --- | --- | --- |
| CADD | 0.00 | 6.7969 | 77.27 | 63.49 | 40.35 | 59.65 | 20.00 | 80.00 | 69.16 | 0.4020 | 0.6814 | 0.5771 | NA | SM |
| CScape | 3.74 | 2.5036 | 55.22 | 55.56 | 30.19 | 69.81 | 60.00 | 40.00 | 55.34 | 0.1028 | 0.4898 | 0.5298 | NA | SM |
| DANN | 0.00 | 11.4002 | **88.89** | 53.93 | 71.93 | 28.07 | **4.00** | **96.00** | 59.81 | 0.3210 | 0.6018 | 0.5879 | NA | SM |
| DIVAN_REGION | 0.00 | 1.8501 | 56.84 | **75.00** | **5.26** | **94.74** | 82.00 | 18.00 | 58.88 | 0.2014 | 0.4351 | NA | 0.5077 | SM |
| DIVAN_TSS | 0.00 | 1.8426 | 55.00 | 71.43 | **3.51** | **96.49** | 90.00 | 10.00 | 56.07 | 0.1310 | 0.3816 | NA | 0.5205 | SM |
| FATHMM-MKL | 0.00 | 7.5595 | 77.27 | 63.49 | 40.35 | 59.65 | 20.00 | 80.00 | 69.16 | 0.4020 | 0.6411 | 0.5502 | 0.5077 | SM |
| FATHMM-XF | 3.74 | 16.2483 | **100.00** | 52.63 | 84.91 | 15.09 | **0.00** | **100.00** | 56.31 | 0.2819 | 0.5270 | 0.5704 | NA | SM |
| FIRE | 0.00 | 3.0552 | 64.47 | **74.19** | **14.04** | **85.96** | 54.00 | 46.00 | 67.29 | 0.3516 | 0.6214 | 0.5250 | 0.5307 | SM |
| ncER | 3.74 | 8.8669 | 84.85 | 61.43 | 49.09 | 50.91 | 10.42 | 89.58 | 68.93 | 0.4329 | 0.7045 | 0.6169 | 0.5396 | SM |
| PAFA | 0.93 | 9.4311 | 67.65 | 54.17 | 58.93 | 41.07 | 22.00 | 78.00 | 58.49 | 0.2040 | 0.5248 | 0.5165 | NA | SM |
| regBase_CAN | 0.00 | 5.5775 | 75.47 | 68.52 | 29.82 | 70.18 | 26.00 | 74.00 | 71.96 | 0.4408 | **0.7775** | 0.5250 | **0.6281** | SM |
| regBase_PAT | 0.00 | 5.3423 | 66.15 | 66.67 | 24.56 | 75.44 | 44.00 | 56.00 | 66.36 | 0.3212 | 0.6926 | 0.6185 | NA | SM |
| regBase_REG | 0.00 | 13.2127 | 81.08 | 61.43 | 47.37 | 52.63 | 14.00 | 86.00 | 68.22 | 0.4052 | 0.7032 | 0.5430 | 0.5230 | SM |
| ReMM | 0.00 | 9.3627 | 75.00 | 61.90 | 42.11 | 57.89 | 22.00 | 78.00 | 67.29 | 0.3640 | 0.7049 | 0.6185 | 0.5018 | SM |
| CDTS | 19.63 | 19.5957 | **100.00** | 61.76 | 59.09 | 40.91 | **0.00** | **100.00** | 69.77 | **0.5027** | 0.6894 | **0.6970** | NA | UM |
| DVAR | 0.00 | 11.7224 | 82.98 | 70.00 | 31.58 | 68.42 | 16.00 | 84.00 | **75.70** | **0.5270** | **0.7970** | **0.6293** | **0.5989** | UM |
| Eigen | 3.74 | 7.8175 | 70.59 | 67.31 | 32.08 | 67.92 | 30.00 | 70.00 | 68.93 | 0.3791 | 0.6743 | **0.6284** | NA | UM |
| Eigen_PC | 3.74 | 12.5576 | 72.34 | 66.07 | 35.85 | 64.15 | 26.00 | 74.00 | 68.93 | 0.3828 | 0.7238 | 0.5491 | NA | UM |
| GenoCanyon | 0.00 | 11.4839 | 79.59 | 68.97 | 31.58 | 68.42 | 20.00 | 80.00 | **73.83** | 0.4849 | 0.7454 | 0.5358 | 0.5485 | UM |
| Orion | 2.80 | 7.5773 | 62.00 | 57.41 | 42.59 | 57.41 | 38.00 | 62.00 | 59.62 | 0.1941 | 0.5641 | 0.5005 | 0.5077 | UM |
| fitCons | 0.93 | 10.3880 | 73.81 | 60.94 | 44.64 | 55.36 | 22.00 | 78.00 | 66.04 | 0.3404 | 0.6159 | 0.5147 | NA | SSM |
| FitCons2 | 0.93 | 11.4394 | 85.37 | 67.69 | 37.50 | 62.50 | 12.00 | 88.00 | **74.53** | **0.5176** | **0.7721** | 0.5513 | **0.5664** | SSM |
| FunSeq2 | 2.80 | 6.6337 | 67.92 | 64.71 | 33.33 | 66.67 | 34.00 | 66.00 | 66.35 | 0.3265 | 0.6811 | 0.5328 | NA | SSM |
| LINSIGHT | 2.80 | 8.0487 | 69.35 | **73.81** | 20.37 | 79.63 | 38.00 | 62.00 | 71.15 | 0.4239 | 0.7448 | 0.5537 | 0.5442 | SSM |

*Note:* Best-threshold, the threshold corresponding to the best sum of sensitivity and specificity; PPV, positive predictive value; NPV, negative predictive value; FPR, false positive rate; FNR, false negative rate; MCC, mathew correlation coefficient; AUC, area under the curve; hspr-threshold, high-specificity regional threshold; hspr-AUC, high-specificity regional area under the curve; hser-threshold, high-sensitivity regional threshold ;hser-AUC, high-sensitivity regional area under the curve; NA, not available. Top three methods of every measure are represented by bold text.

**Table S8.** Performance of 24 methods based on 4 benchmark datasets.

| Method | Germline | Somatic | eQTL | GWAS |
| --- | --- | --- | --- | --- |
| CADD | I | III | II | I |
| CDTS | III | II | III | I |
| CScape | II | II | III | III |
| DANN | II | III | III | II |
| DIVAN_REGION | III | III | I | I |
| DIVAN_TSS | III | III | I | I |
| DVAR | II | I | II | II |
| Eigen | I | II | I | II |
| Eigen_PC | III | I | I | II |
| FATHMM-MKL | I | II | II | III |
| FATHMM-XF | I | III | III | III |
| FIRE | III | II | I | II |
| fitCons | III | II | II | II |
| FitCons2 | II | I | I | III |
| FunSeq2 | II | I | II | II |
| GenoCanyon | III | I | II | II |
| LINSIGHT | I | I | I | I |
| ncER | I | I | II | III |
| Orion | III | III | III | I |
| PAFA | II | III | II | I |
| regBase_CAN | II | I | III | III |
| regBase_PAT | I | II | III | III |
| regBase_REG | II | II | I | I |
| ReMM | I | III | III | III |

*Note:* We divided 24 methods into three groups (I, II and III) based on the rank of AUC values and every group contained 8 methods. eQTL, expression quantitative trait locus; GWAS,

**Table S9.** Matched configurations of vSampler.

| Matched criteria | configuration |
| --- | --- |
| Minor allele frequency deviation | -0.01, 0.01 |
| Distance to closest transcription start site deviation | -5KB, 5KB |
| Gene density in distance | 100KB |
| Gene density deviation | -5, 5 |
| Linkage disequilibrium threshold | $r^2 > 0.8$ |
| Number of variants in linkage disequilibrium deviation | -5, 5 |

